# Large-scale single-cell profiling of stem cells uncovers redundant regulators of shoot development and yield trait variation

**DOI:** 10.1101/2024.03.04.583414

**Authors:** Xiaosa Xu, Michael Passalacqua, Brian Rice, Edgar Demesa-Arevalo, Mikiko Kojima, Yumiko Takebayashi, Benjamin Harris, Hitoshi Sakakibara, Andrea Gallavotti, Jesse Gillis, David Jackson

## Abstract

Stem cells in plant shoots are a rare population of cells that produce leaves, fruits and seeds, vital sources for food and bioethanol. Uncovering regulators expressed in these stem cells will inform crop engineering to boost productivity. Single-cell analysis is a powerful tool for identifying regulators expressed in specific groups of cells. However, accessing plant shoot stem cells is challenging. Recent single-cell analyses of plant shoots have not captured these cells, and failed to detect stem cell regulators like *CLAVATA3* and *WUSCHEL*. In this study, we finely dissected stem cell-enriched shoot tissues from both maize and arabidopsis for single-cell RNA-seq profiling. We optimized protocols to efficiently recover thousands of *CLAVATA3* and *WUSCHEL* expressed cells. A cross-species comparison identified conserved stem cell regulators between maize and arabidopsis. We also performed single-cell RNA-seq on maize stem cell overproliferation mutants to find additional candidate regulators. Expression of candidate stem cell genes was validated using spatial transcriptomics, and we functionally confirmed roles in shoot development. These candidates include a family of ribosome-associated RNA-binding proteins, and two families of sugar kinase genes related to hypoxia signaling and cytokinin hormone homeostasis. These large-scale single-cell profiling of stem cells provide a resource for mining stem cell regulators, which show significant association with yield traits. Overall, our discoveries advance the understanding of shoot development and open avenues for manipulating diverse crops to enhance food and energy security.

## INTRODUCTION

Stem cells in shoot apical meristems shape plant architecture and produce fruits and seeds, which are crucial for food and carbon neutral biofuel production (Greb and Lohmann, 2016; Lindsay et al., 2024). Understanding the control of these stem cells, including their maintenance and proliferation, requires the identification and functional characterization of master stem cell regulators within complex regulatory networks. Genetics and bulk gene expression approaches have identified and localized some of these regulators (Kitagawa and Jackson, 2019; Lindsay et al., 2024). This has led to the identification of specific cell types and domains, including the central zone which harbors stem cells, the organizing center which specifies the stem cells, and the peripheral zone which initiates lateral organ primordia (Fuchs and Lohmann, 2020; Hong and Fletcher, 2023; Karami et al., 2023; Uchida and Torii, 2019; Wang et al., 2020; Zhang et al., 2023). These cell types and developmental domains are highly conserved in both vegetative and inflorescence shoot meristems, and across different plant species (Greb and Lohmann, 2016). However, a comprehensive cross-species identification of conserved shoot meristem regulators is lacking.

Among known shoot stem cell regulators, the signaling peptide CLAVATA3 (CLV3) and homeodomain transcription factor WUSCHEL (WUS) form a negative feedback-loop to coordinate stem cell proliferation and differentiation (Hirakawa, 2021; Somssich et al., 2016). In arabidopsis, *CLV3* is expressed in stem cells to restrict the expression of WUS to the organizing center, and WUS protein moves from the organizing center to stem cells to activate *CLV3* expression (Daum et al., 2014; Yadav et al., 2011). The identification of shoot stem cell regulators is critical to understanding the mechanisms of stem cell self-renewal, proliferation, and differentiation, and they can be manipulated by genome editing to bolster crop yield (Lindsay et al., 2024; L. Liu et al., 2021b; Rodríguez-Leal et al., 2017; Rodriguez-Leal et al., 2019; Wang et al., 2021).

Transcriptomic approaches leveraging Fluorescence-Activated Cell Sorting (FACS) or Translating Ribosome Affinity Purification (TRAP) have identified arabidopsis genes that are highly expressed in the *CLV3* and *WUS*-expressing cells (Tian et al., 2019; Yadav et al., 2009). However, this approach relied on transgenic lines and low-throughput isolation protocols that cannot be easily applied to crop plants. In recent years, single-cell transcriptomic analysis has been extensively used to profile shoot meristems in arabidopsis (Neumann et al., 2022; Zhang et al., 2021b), maize (Satterlee et al., 2020; Xu et al., 2021), rice (Zong et al., 2022), and Populus (Conde et al., 2022). However, these studies failed to recover cells expressing *CLV3* and/ or *WUS,* presumably due to their scarcity, inaccessibility, fragility, or low expression level of these markers.

In this study, we overcame this challenge by finely dissecting stem cell-enriched tissues from inflorescence apices and optimizing the cell isolation protocol to preserve rare stem cells, followed by single-cell RNA-sequencing. We recovered ∼5,000 *CLV3* and ∼1,000 *WUS*-expressing cells and identified hundreds of stem cell markers in both maize and arabidopsis. A single-cell cross-species analysis identified conserved stem cell markers and cell types that we validated using high-resolution spatial transcriptomics. We functionally characterized a family of conserved stem cell markers orthologous to human SERPINE1 mRNA binding protein (SERBP1), a key ribosome associated factor in humans. Furthermore, we profiled maize stem cell overproliferation mutants and uncovered two families of sugar kinase genes that are candidate targets of WUS. One family of these genes is controlled by hypoxia signaling, while the other family functions in cytokinin hormone homeostasis in stem cells. Our large-scale single-cell profiling is a valuable resource for mining stem cell regulators and understanding hidden regulatory mechanisms in plant shoot development. We observed a high association between these stem cell regulators and maize ear yield traits, which informed future crop engineering and improvement. Our analyses will inform stem cell studies in other crops, and provide avenues for food and energy security.

## RESULTS

### Recovery of rare stem cells in single-cell profiling of maize and arabidopsis inflorescences

To recover stem cells that were largely missed in previous shoot single-cell studies (Conde et al., 2022; Neumann et al., 2022; Satterlee et al., 2020; Xu et al., 2021; Zhang et al., 2021b; Zong et al., 2022), we finely dissected the tips of developing maize ear primordia (wild type B73), comprising the inflorescence meristem and early stage spikelet pair meristems (Figure 1A). Additionally, we optimized our cell sorting protocol (Xu et al., 2021) using OptiPrep Gradient Medium (Sigma-Aldrich, OptiPrep™ Application Sheet C18), or by modifying the cell isolation steps based on (Satterlee et al., 2020) (see STAR methods for details). We loaded the protoplasts into a 10x Genomics Chromium Controller, followed by library construction and sequencing. We mapped the sequencing reads to the maize v3 reference genome and clustered the cells with Seurat (Hao et al., 2021). We successfully recovered stem cells in all replicates of both approaches. Thus, we integrated all the datasets using Seurat for single-cell clustering analysis (Hao et al., 2021). In total, we recovered 1,208 stem cells marked by the maize *CLV3* ortholog, *ZmCLAVATA3/EMBRYO SURROUNDING REGION-RELATED7* (*ZmCLE7*) and 112 cells marked by maize *WUSCHEL1* (*ZmWUS1*) (Figures 1B-1C). We also recovered all known cell or domain types with a total of 11,554 cells, including epidermis, meristem base, adaxial meristem periphery, determinate lateral organ cortex, and pith, and identified hundreds of cell type-specific markers (Figures 1B and S1A-B; Table S1). We also separated the inflorescence meristem (cluster 3) and early stage spikelet pair meristems (cluster 5) based on the differential expression of axillary meristem marker genes, *BARREN INFLORESCENCE1* and *BARREN INFLORESCENCE4* (*BIF1* and *BIF4*) (Galli et al., 2015) (Figures 1A-C and S1C).

**Figure 1:**
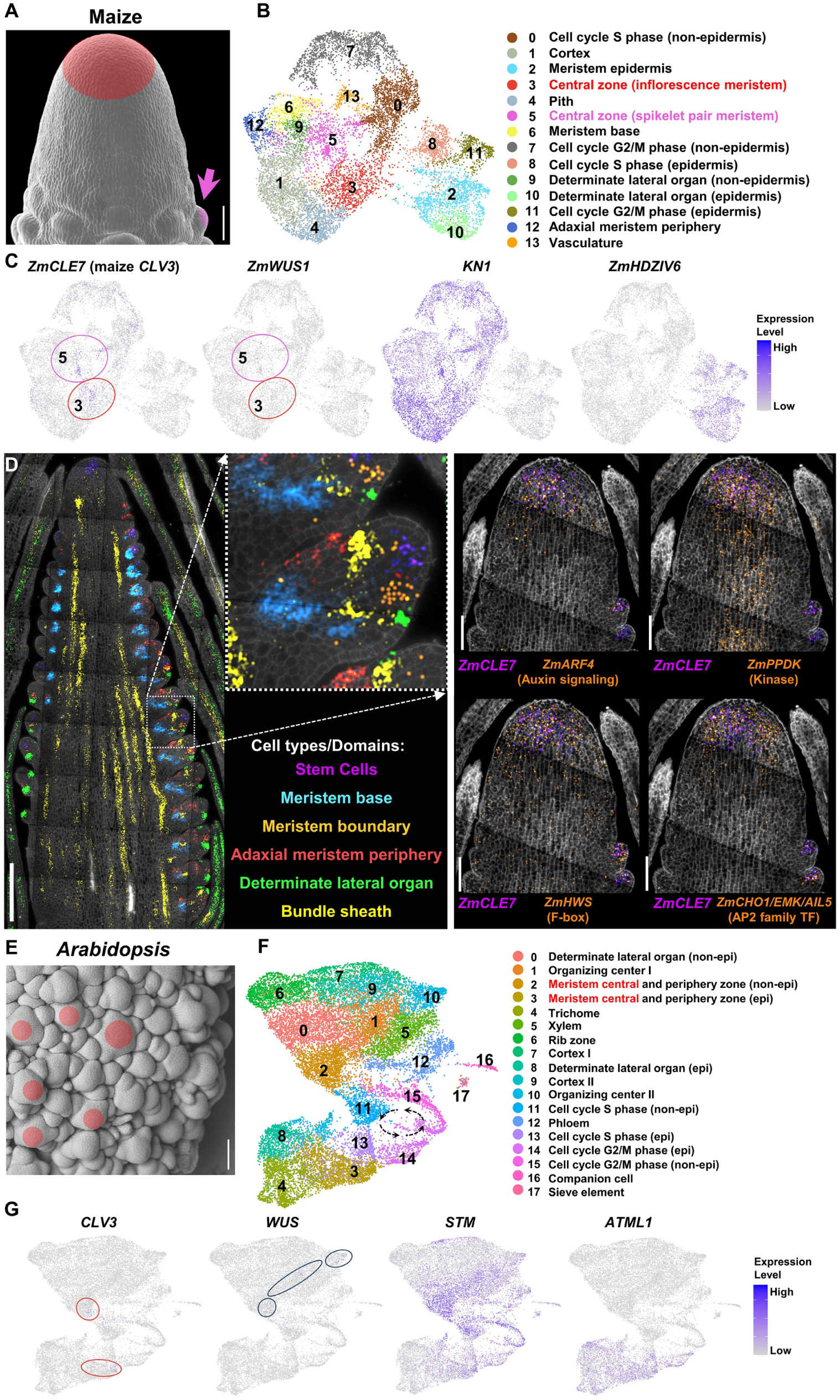
Single-cell RNA-seq analysis of inflorescence shoot stem cells in maize and arabidopsis. (A) Scanning electron microscopy image of a finely dissected maize B73 ear tip (scale bar = 100µm). Stem cells are shaded in inflorescence meristem (red) and spikelet pair meristem (pink). (B) 14 distinct single-cell RNA-seq clusters of maize B73 ear tip. (C) Uniform Manifold Approximation and Projection (UMAP) plots of *ZmCLE7*, ZmWUS1, *KNOTTED1* (*KN1*), and *ZmHOMEODOMAIN LEUCINE ZIPPER IV6* (*ZmHDZIV6*) reveal two central zone clusters harboring stem cells (cluster 3, inflorescence meristems; cluster 5, early stage spikelet pair meristems). Color scale indicates normalized expression level. (D) Left panel: spatial transcriptomic analysis showing distinct cell types of maize developing ear marked by expression of *ZmCLE7* (stem cells), *BRANCHED SILKLESS1* (*BD1*) (meristem boundaries), *Zm1-AMINOCYCLOPROPANE-1-CARBOXYLATE OXIDASE2* (*ZmACO2*) (adaxial meristem periphery), *RAMOSA* (*RA3*) (meristem base), *ZmYABBY14* (*ZmYAB14*) (determinate lateral organ), and *ZmSHORT-ROOT* (*ZmSHR1*) (bundle sheath), scale bar = 500µm; Right panel: spatial gene expressional validation of *ZmCLE7* (magenta) and highly co-expressed markers (gold), including *ZmARF4*, *ZmPPDK*, *ZmHWS*, and *ZmCHO1/EMK/AIL5*, in stem cells, scale bar = 100µm. (E) Scanning electron microscopy image of dissected arabidopsis *ap1;cal* mutant inflorescence (scale bar = 100µm). Stem cells are shaded for a few representative meristems (red). (F) 18 distinct single-cell RNA-seq clusters of arabidopsis *ap1;cal* mutant. (G) UMAP plots of *CLV3*, *WUS*, *SHOOT MERISTEMLESS* (*STM*), and arabidopsis *THALIANA MERISTEM LAYER 1* (*ATML1*) revealing stem cells in a part of cluster 2 (non-epidermis) and cluster 3 (epidermis). Epi= epidermis. Color scale indicates normalized expression level.

The successful recovery of *ZmCLE7* marked stem cells enabled us to identify novel stem cell regulators by performing a co-expression analysis. We identified the top 300 *ZmCLE7* co-expressed genes, which we considered highly confident stem cell marker candidates (see STAR Methods; Table S2). Among these candidates were known shoot stem cell regulators, *FASCIATED EAR4* (*FEA4*) (Pautler et al., 2015), *ZmLONELY GUY7* (*ZmLOG7*) and *ZmFON2-LIKE CLE PROTEIN1* (*ZmFCP1*) (Knauer et al., 2019). Gene Ontology (GO) enrichment analyses of the highly co-expressed markers identified significant enrichment for primary shoot apical meristem specification, flower development, and phyllotactic patterning. We also found significant enrichment in hormone signaling (auxin, cytokinin, and abscisic acid), biosynthesis and/or homeostasis (auxin, gibberellin, cytokinin, and ethylene), catabolic process (salicylic acid, abscisic acid, and gibberellin) and transport (auxin). Translation, gene expression, protein metabolic process and modification, RNA binding, ribosomal assembly and mitochondrial complex were also enriched (Table S2). To validate these markers, we first performed *in situ* hybridization for *ZmWRKY14* (Tang et al., 2021) and *ZmPHABULOSA489* (*ZmPHB489*) (Du et al., 2021) transcription factors and a homolog of arabidopsis F-box gene *HAWAIIAN SKIRT* (*ZmHWS*) (Zhang et al., 2017). Expression of these genes was indeed enriched in stem cells (Figure S1D). To increase the validation throughput, we next used a spatial transcriptomics approach based on Resolve Biosciences’ Molecular Cartography. We first confirmed distinct cell types predicted by previous single-cell RNA-seq (Xu et al., 2021) in a spatial context (Figure 1D). Next, we validated the expression of newly identified stem cell marker candidates, such as *ZmARF4,* a homolog of arabidopsis auxin signaling regulator *MONOPTEROS* (Galli et al., 2015; Schlereth et al., 2010), *ZmPPDK,* a homolog of arabidopsis *PYRUVATE ORTHOPHOSPHATE DIKINASE* (Moons et al., 1998; Parsley and Hibberd, 2006), and *ZmCHO1/EMK/AIL5,* a homolog of arabidopsis AP2 family transcription factor, *CHOTTO1/EMBRYOMAKER/AINTEGUMENTA-like 5* (Nole-Wilson et al., 2005), and the F-box gene *ZmHWS* (Zhang et al., 2017) (Figure 1D).

To capture gene expression dynamics in the developmental progression from inflorescence meristem to spikelet pair meristem (Fig.1A), we next used the Slingshot algorithm for pseudotime inference (Figure S2). For the inflorescence meristem, we sub-clustered central zone stem cell cluster 3 and its adjacent differentiating ground tissue clusters, cortex (cluster 1), and pith (cluster 4). After calculating the pseudotime values and mapping the trajectory to the cell clusters, we captured a branching trajectory from central zone stem cell cluster 3 to the two differentiating ground tissue clusters, 1 and 4 (Figures S2A-D). Projection of *ZmCLE7* expression to the two lineages revealed high expression at the beginning and lower expression towards the end, matching the pattern shown by *in situ* hybridization (Figures 2E-2F). Similarly, for early stage spikelet pair meristems, we sub-clustered central zone stem cell cluster 5 and the differentiating clusters, including meristem base cluster 6, adaxial meristem periphery clusters 12, determinate lateral organ cluster 9, and vasculature cluster 13. We again observed branching trajectories (Figure S2G-H) and a decrease in *ZmCLE7* expression along each lineage, as expected (Figures S2I-J).

**Figure 2:**
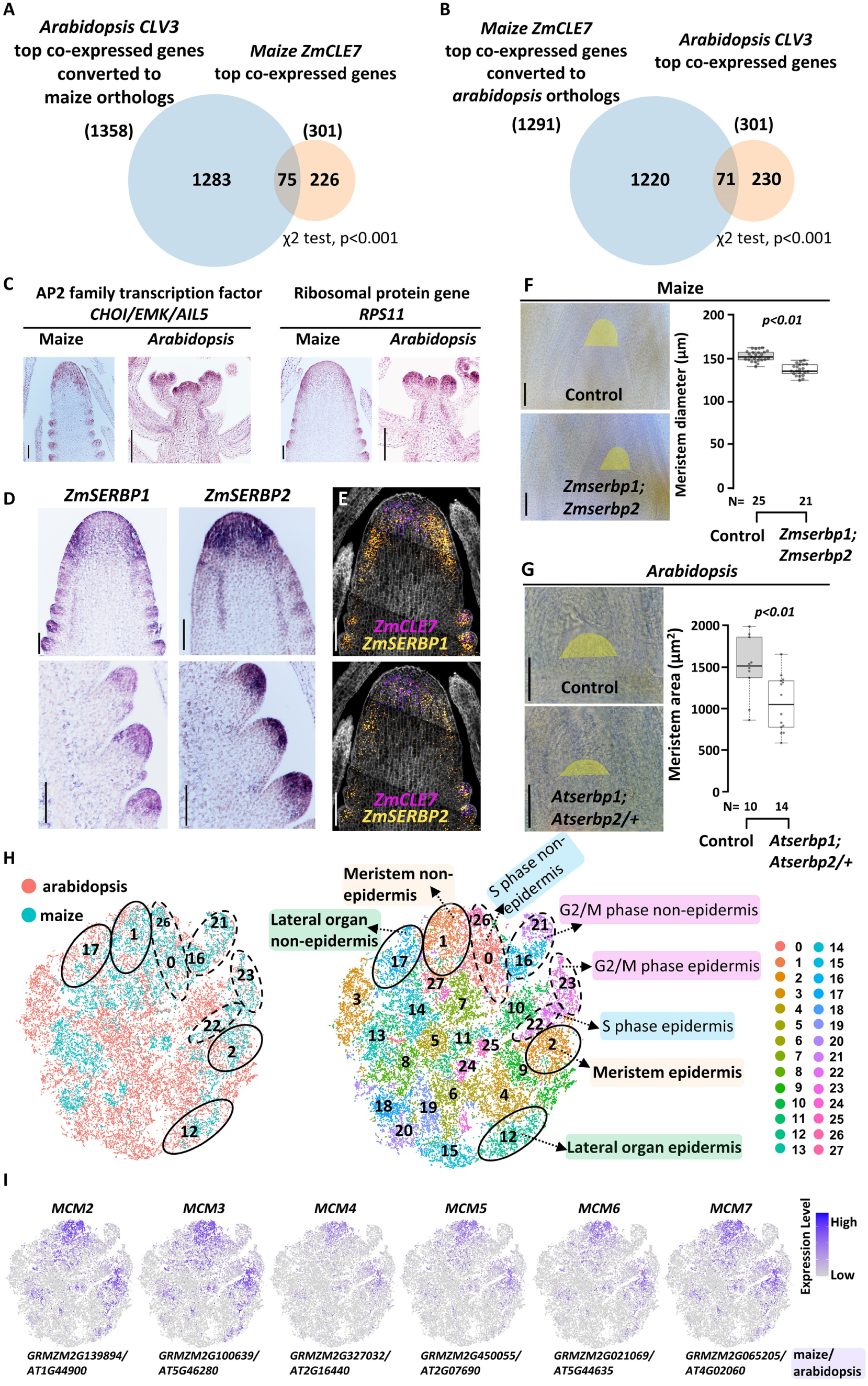
Cross-species comparison of shoot stem cells at single-cell resolution. (A) Top 300 *CLV3* co-expressed genes in arabidopsis converted to 1358 maize orthologs; 75 of them overlap with top 300 *ZmCLE7* co-expressed genes in maize. Overlap is significant, χ2 test, p<0.001. (B) Top 300 *ZmCLE7* co-expressed genes in maize converted to 1291 arabidopsis orthologs, 71 of them overlap with top 300 *CLV3* co-expressed genes in arabidopsis. Overlap is significant, χ2 test, p<0.001. (C) mRNA *in situ* of newly identified stem cell marker genes validates their predicted conserved stem cell expression in both species. Scale bar = 100µm. (D-E) *in situ* hybridization validation of *ZmSERBPs* (D) and spatial transcriptomics (E), scale bar = 100µm. (F) Maize *Zmserbp1;Zmserbp2* double mutants and arabidopsis *Atserbp1;Atserbp2/+* mutants have smaller shoot apical meristems (One-Way ANOVA, p<0.005). Scale bar = 100µm (F) and 50µm (G) respectively. The horizontal line within the box represents the median value. (H) t-Distributed Stochastic Neighbor Embedding (*t*-SNE) plots shows the integration of arabidopsis and maize shoot tip cells (left panel) resulting in 28 cell clusters (right panel). Dotted ovals indicate dividing cell clusters, solid ovals indicate the conserved meristem and determinate lateral organ clusters. (I) *t*-SNE plots of a group of conserved meristem marker genes that belong to *MINICHROMOSOME MAINTENANCE* (*MCM*) family in maize and arabidopsis. The dots represent all cells from maize and arabidopsis in maize and arabidopsis integrated atlas shown in (H, right panel). Color scales indicate normalized expression level.

We next captured rare shoot stem cells in arabidopsis (Neumann et al., 2022; Zhang et al., 2017), using *apetala1*; *cauliflower* (*ap1;cal*) double mutants, with over proliferating inflorescence meristems (Figure 1E) (Kempin et al., 1995). The CLV-WUS network is operational in this mutant background (Yadav et al., 2009). Applying the same optimized protoplasting conditions (see STAR methods), we recovered 392 arabidopsis inflorescence shoot stem cells marked by *CLV3* from epidermis and inner layers (Figures 1E-1G), as well as many other distinct cell types (Figures 1F and S3A-B; Table S1). We also captured 316 *WUS*-expressing cells (Figures 1F-1G) and found that a small number of them overlapped with *CLV3*-expressing cells, as previously reported (Yadav et al., 2009, PNAS). We compared our single-cell RNA-seq with FACS-microarray results (Yadav et al., 2009), using single-cell co-expression analysis of *CLV3* and *FILAMENTOUS FLOWER* (*FIL*) (Table S1). We selected the top 300 highly co-expressed markers for each gene, and found significant overlap (p < 0.001, chi-square test) with the markers identified by FACS-microarray (Figure S3C) (Yadav et al., 2009). GO analysis of the top 300 *CLV3* co-expressed markers identified gene expression regulation, RNA binding, ribosomal assembly, and mitochondrial complex categories. We also found strong enrichment of meristem development, stem cell population maintenance, chromatin organization, and cytokinin hormone metabolic process categories (Table S2). These categories were similar to our analysis of *ZmCLE7* co-expressed markers in maize (Table S2).

### A cross-species analysis of inflorescence stem cells at single-cell resolution

Previous studies identified a limited number of evolutionarily conserved stem cell regulators (Chen et al., 2021; Lohmann et al., 2011; Mayer et al., 1998; Pautler et al., 2015; Rodriguez-Leal et al., 2019). Our capture of maize and arabidopsis stem cells allowed us to perform a more comprehensive single-cell cross-species analysis at both stem cell marker and cell type levels. At the stem cell marker level, we first identified the maize orthologs of the top 300 *CLV3* co-expressed markers in arabidopsis (STAR methods) and found that 75 of them overlapped with the top 300 *ZmCLE7* co-expressed markers (chi-square test, p<0.001, Figure 2A, Table S2). We also identified 71 orthologs of *ZmCLE7* co-expressed markers from maize that significantly overlapped with *CLV3* co-expressed markers (chi-square test, p<0.001, Figure 2B, Table S2). Among these highly conserved markers, we found the conserved stem cell marker *FEA4/ PERIANTHIA* (*PAN*) bZIP transcription factors enriched in both maize (*FEA4*) and arabidopsis (*PAN*) inflorescence stem cells (Table S2) (Lohmann et al., 2011; Pautler et al., 2015). We also validated expression of a newly identified conserved stem cell marker, the AP2 family transcription factor *CHO1/EMK/AIL5* (Figures 2C and 1D).

We next functionally validated a group of conserved stem cell marker genes that have not been studied in plants but are orthologous to human *SERPINE1 mRNA BINDING PROTEIN* (*ZmSERBP*) (Heaton et al., 2001). Two out of three members of the maize *ZmSERBP* family, *ZmSERBP1* and *ZmSERBP2*, were identified as conserved stem cell markers, and validated using mRNA *in situ* and spatial transcriptomics (Figures 2D-2E). We then generated Clustered Regularly Interspaced Short Palindromic Repeats (CRISPR)-Cas9 mutants for both *ZmSERBP* genes (Figure S4) and found that the double mutant had smaller shoot apical meristems (Figure 2F). We also characterized arabidopsis *SERBP* T-DNA insertion mutants, but failed to recover *Atserbp1;Atserbp2* double mutants, possibly due to lethality. However, plants homozygous for *Atserbp1* and heterozygous for *Atserbp2* also had smaller shoot apical meristems (Fig2G). These results indicate a conserved function of SERBPs in regulating shoot meristem development in maize and arabidopsis. Human SERBP1 regulates translation by associating with ribosomes (Baudin et al., 2021; Brown et al., 2018; Heaton et al., 2001; Muto et al., 2018). Interestingly, ∼70% of the stem cell markers conserved between maize and arabidopsis encoded ribosomal proteins (Table S2), and mRNA *in situ* hybridization validated their enriched expression in maize and arabidopsis stem cells (Figures 2C and S5A). These results support a hypothetical conserved stem cell role for specialized ribosomes (Barna et al., 2022; Genuth et al., 2022; Genuth and Barna, 2018), as well as the associated SERBPs (Li et al., 2016) across kingdoms.

Next, we asked if shoot meristem cell-types were conserved between maize and arabidopsis. Such analyses usually rely on 1-to-1 orthologs between species (Zhang et al., 2021a). However, 1- to-1 orthologs are rare in plants due to whole genome and tandem gene duplications, making integration difficult. We therefore used RNA-seq data from both species to identify pairs of genes with highly similar co-expression profiles to integrate cell-type-specific data. We used a computational algorithm, Expression Proxies In Plants Help Integrate Transcribed Expression in Single-cell (EPIPHITES) (Passalacqua and Gillis, 2023) to identify such gene pairs (https://gillislab.shinyapps.io/epiphites/). We found 6,075 co-expressed gene pairs between maize and arabidopsis, and used them to integrate our single-cell datasets (Figures 1B and 1F) and identified 28 cell clusters by Seurat CCA (Figure 2H, see STAR methods). By projecting the integrated clusters back to arabidopsis cluster annotations (Figure S5B), we found that meristem (clusters 1 and 2) and determinate lateral organ (clusters 17 and 12) cells were highly conserved across the two species (Figures 2H). This was supported by marker annotations, including *ERECTA-like1*, functioning in meristems (Shpak et al., 2005) and *ETHYLENE AND SALT INDUCIBLE 3* (*ESE3*), involved in leaf organ identity (Lumba et al., 2012) (Figure S5C). We also found strong similarity between the two species for epidermal (clusters 2, 4, 9, 10, 12, 22, and 23) and dividing cell clusters (G2/M phase, clusters 16, 21, and 23; S phase, clusters 0, 22, and 26) (Figures 2H-2I and S5D). We next compiled lists of highly conserved cross-species marker genes enriched in the conserved meristem and determinate lateral organ cell types (Table S3). GO analysis of conserved marker genes found over-representation of meristem development, reproductive structure development, signal transduction, cell communication and response to hormones categories (Table S3). Metabolic processes, DNA replication and repair, translation and ribosomal assembly were also enriched (Table S3). Strikingly, we found a group of conserved meristem markers that belong to a family of *MINICHROMOSOME MAINTENANCE* (*MCM*) proteins (Figure 2I), which have critical roles in ear (cob) development in maize (Dresselhaus et al., 2006), and function in embryo and root meristem in arabidopsis (Ni et al., 2009), indicating these genes may play a conserved role in different tissues across species.

### Single-cell analysis of stem cell over-proliferation mutants

Our success in stem cell profiling motivated us to next investigate stem cell regulatory mutants at the single-cell level. We profiled a *fasciated ear3;Zmcle7* (*fea3;Zmcle7*) double mutant (Je et al., 2016; Rodriguez-Leal et al., 2019) which has very severe over-proliferation of ear tip stem cells (Figures S4A and 3A; see STAR methods), and the dominant *Barren inflorescence3* (*Bif3*) mutant (Chen et al., 2021), in which stem cells over-proliferate due to *ZmWUS1* overexpression. A single-cell gene expression atlas of *fea3;Zmcle7* ear tips, harboring inflorescence meristem and early stage spikelet pair meristems identified distinct cell types, similar to wild type (Figures S6A-S6B). We recovered 3,017 *ZmCLE7* and 145 *ZmWUS1* expressing cells. Next, we integrated *fea3;Zmcle7* and wild type datasets using Seurat, and identified obvious stem cell clusters in both inflorescence and early stage spikelet pair meristems (Figures 3A and S7). As expected, the number of stem cells was significantly higher in the *fea3;Zmcle7* mutants (Figures 3A-B), and was validated by mRNA *in situ* of *ZmCLE7* (Figure 3B), whose expression per-cell was also significantly higher in *fea3;Zmcle7* mutants (Figure 3C). We also profiled *Bif3* homozygous mutant ear tips, (Figure S8A), and again identified stem cell clusters from both inflorescence and spikelet pair meristems, with a total of 315 *ZmCLE7* and 450 *ZmWUS1* expressing cells (Figure S8B). After integration with wild type, we again identified two stem cells clusters corresponding to both types of meristems (Figures 3D and S9). As expected, *ZmWUS1* expression expanded in *Bif3* mutants (Figure 3E) (Chen et al., 2021), and was upregulated in the mutant stem cells (Figure 3F).

**Figure 3:**
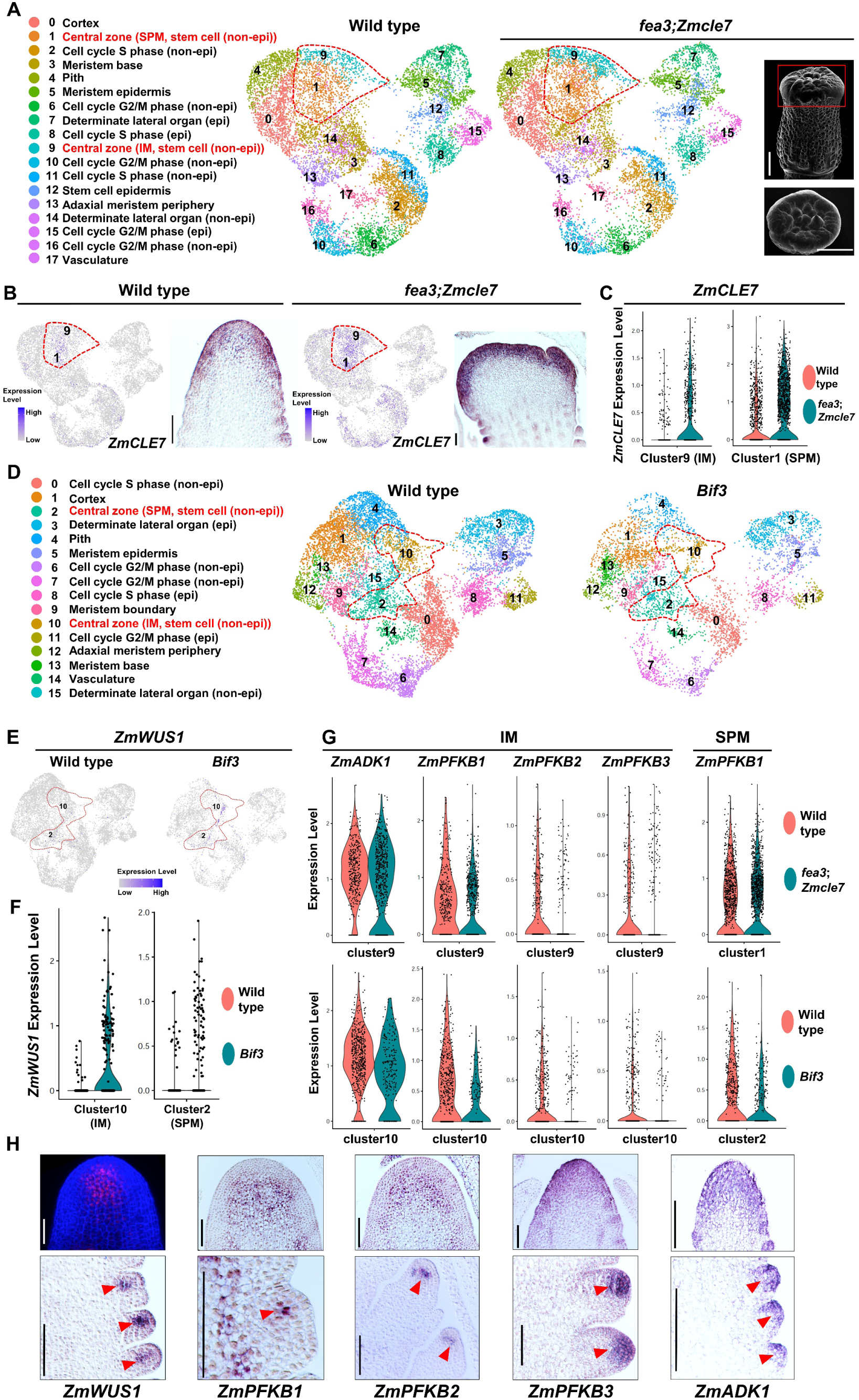
single-cell RNA-seq analysis of stem cell proliferation mutants. (A) Integration and annotation of wild type and *fea3;Zmcle7* ear tip cells. The non-epidermal central zone stem cells in inflorescence meristem (cluster 9) and early stage spikelet pair meristem (cluster 1) are highlighted with red dotted lines. *fea3;Zmcle7* ear tips were finely dissected as marked by the red box of scanning electron microscopy image (upper panel; top view in lower panel (scale bar = 1mm). (B) UMAP plots and mRNA *in situ* hybridization of *ZmCLE7* in wild type and *fea3;Zmcle7* double mutants. Color scale in UMAP plots indicates normalized expression level. (C) Violin plots show upregulated *ZmCLE7* expression in the stem cells of *fea3;Zmcle7* double mutants. (D) UMAP plots of wild type B73 and *Bif3* mutant ear tip cells with annotation after integration. (E) UMAP plots showing higher expression of *ZmWUS1* in the *Bif3* central zone clusters in inflorescence meristem (cluster 10) and spikelet pair meristem (cluster 2). Color scale in UMAP plots indicate normalized expression level. (F) Violin plots of *ZmWUS1* show upregulated expression in *Bif3* mutant stem cells. (G) Violin plots show downregulation of *ZmADK* and *ZmPFKB* genes in stem cells of *fea3;Zmcle7* and *Bif3* mutants. (H) *pZmWUS1*-mRFP reporter line (Je et al., 2016) and mRNA *in situ* hybridization of *ZmWUS1, ZmPFKBs,* and *ZmADK1*. Scale bar = 100µm.

Next, we asked which genes were differentially expressed in *fea3;Zmcle7* or *Bif3* mutant stem cells and might underly their stem cell over-proliferation phenotypes (see STAR methods). To narrow down high confidence candidates, we performed stringent *Bonferroni* correction to adjust combined probability (p)-values from comparisons between both mutants and wild type (Table S4). By setting the Bonferroni adjusted p-value threshold to < 0.05, we identified 4,095 and 3,806 differentially expressed genes in inflorescence meristem and spikelet pair meristem stem cells, respectively. Among these candidates, we found that *PHOSPHOFRUCTOKINASE B (PFKB)-TYPE* carbohydrate kinase genes (Schroeder et al., 2018), and related *ZmADENOSINE KINASE* (*ZmADK*) genes (Schoor et al., 2011) were differentially expressed in the *fea3;Zmcle7* and *Bif3* mutants stem cells (Figures 3G; Table S4).

*ZmPFKB* and *ZmADK* genes belong to a large family of predicted sugar metabolic genes (Figure S10). Using mRNA *in situ* hybridization, we found meristem enriched expression of *ZmPFKB1* and *ZmPFKB2* that largely overlapped with *ZmWUS1* (Figure 3H). This suggests that *ZmPFKB* genes might be regulated by ZmWUS1. Indeed, a recent Cleavage Under Targets & Tagmentation (tsCUT&Tag) analysis found several *ZmPFKB* genes as direct binding targets of ZmWUS1 (Figure 4A) (Dong et al., 2022). Similarly, *PFKB* orthologs *AT4G28706*, *AT5G43910*, and *AT5G58730* were identified as WUS targets in arabidopsis (Ma et al., 2019). Since several *ZmPFKB* genes were expressed in stem cells, we reasoned that they are likely to act redundantly, and made knock outs using multiplex CRISPR-Cas9 (Figure S11). We found that triple *Zmpfkb* mutants (*Zmpfkb1;2;3*) had enlarged shoot apical meristems (Figure 4B). A *PFKB* gene is regulated by hypoxia in bacteria (Shi et al., 2010), and hypoxia promotes stem cell proliferation in humans (Li et al., 2021). The stem cell niche in arabidopsis is also a hypoxic zone, and a Leucine Zipper gene, *LITTLE ZIPPER 2* (*ZPR2*), is induced under hypoxia (Weits et al., 2019). Maize *LITTLE ZIPPER* homolog *ZmLITTLE ZIPPER2* (*ZmZPR2*) had enriched expression in stem cells (Figure 4C). Since expression of *ZmPFKB* genes was enriched in stem cells, and these genes were differentially expressed in stem cell overproliferation mutants (Figures 3G-H), we hypothesized that *ZmPFKBs* might respond to hypoxia signaling in maize shoot stem cells. To test our hypothesis, we incubated ear primordia in hypoxic conditions (2% O_2_), and indeed found up-regulation of *ZmZPR2* as in arabidopsis (Figure 4D) (Weits et al., 2019). Strikingly, we found significant downregulation of *ZmPFKB1* and *ZmPFKB2* genes under hypoxia, similar to their downregulation observed in stem cell mutants (Figures 4D and 3G). Overall, our single-cell analysis enabled us to uncover a group of ZmWUS1 candidate target *ZmPFKB* genes, that were regulated by hypoxia signaling.

**Figure 4:**
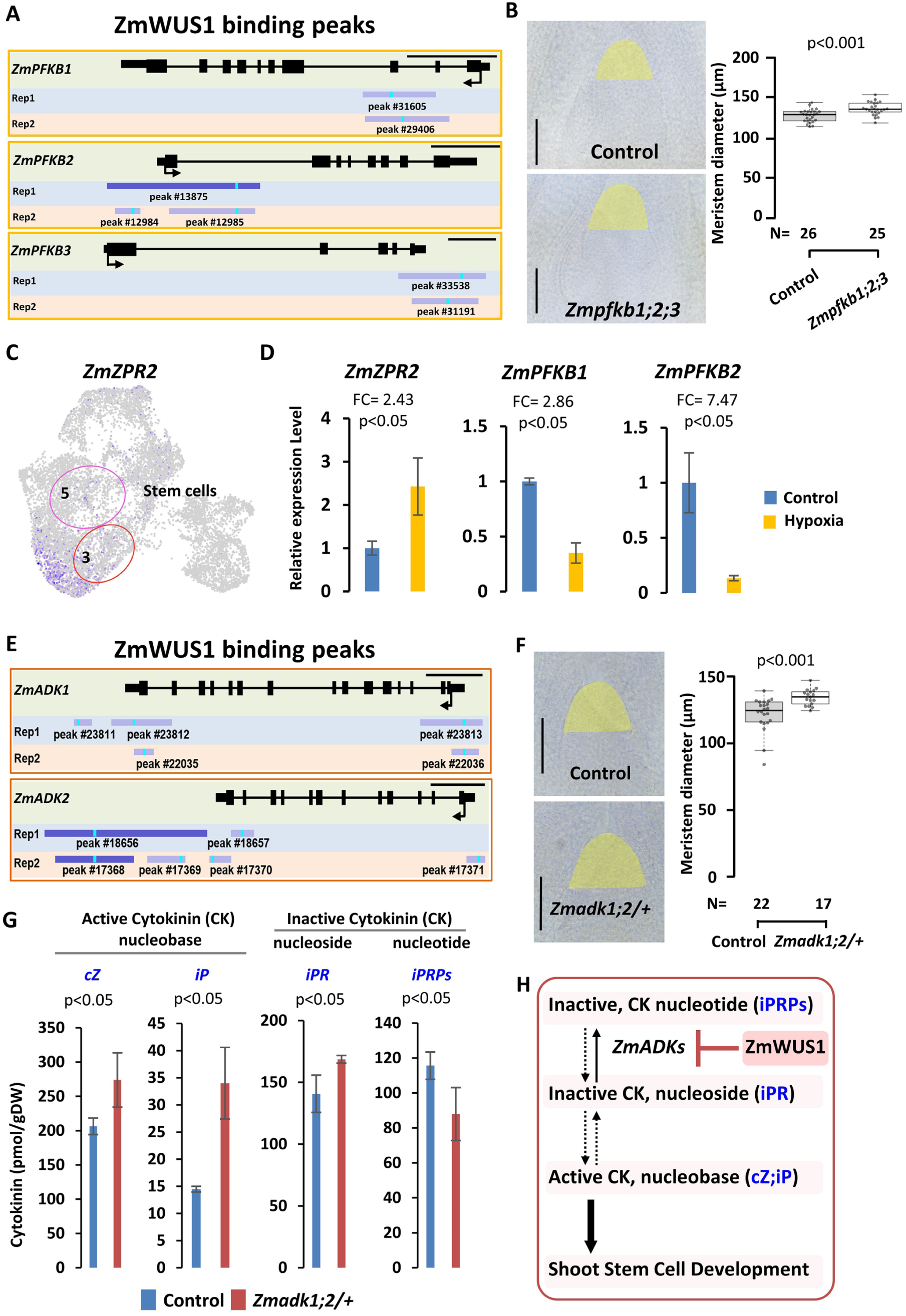
*ZmPFKB* and ZmADK genes control shoot meristem development and are putative ZmWUS1 targets. (A) ZmWUS1 binding peaks in *ZmPFKB* genes from a tsCUT-Tag assay (Dong et al., 2022). Large peaks in dark blue color; small peaks in light purple color; peak summits in bright blue color. (B) Multiplex CRISPR-Cas9 mutants of *Zmpfkb1;2;3* have enlarged shoot apical meristems (One-Way ANOVA, p<0.001). Scale bar = 100µm. The horizontal line within the box represents the median value. (C) UMAP plot of *ZmZPR2* showing expression in the two stem cell clusters (cluster 3, inflorescence meristems; cluster 5, early stage spikelet pair meristems). Color scale indicates normalized expression level. (D) qPCR analysis of *ZmZPR2* and *ZmPFKBs* expression in response to hypoxia. (E) ZmWUS1 binding peaks in *ZmADK* genes from the tsCUT-Tag assay (Dong et al., 2022), large peaks in dark blue color; small peaks in light purple color; peak summits in bright blue color. (F) *Zmadk1;2/+* mutant shoot apical meristems are significantly larger (One-Way ANOVA, p<0.001). Scale bar = 100µm. The horizontal line within the box represents the median. (G) Quantification of different forms of cytokinin (CK). (H) Regulation model of cytokinin homeostasis by *ZmADKs* and ZmWUS1 underlying shoot stem cell development.

We next investigated the spatial expression and function of *ZmADK* genes. *ZmADK* expression was enriched in stem cells in a similar pattern as *ZmCLE7* (Figures 3H and 3B). Given that WUS protein is mobile (Yadav et al., 2011), we hypothesized that *ZmADK* genes might also be regulated by ZmWUS1. Indeed, several *ZmADK* genes are candidate direct binding targets of ZmWUS1 (Figure 4E) (Dong et al., 2022). To investigate *ZmADK* function, we again used multiplex CRISPR-Cas9 (Figures 4F and S11). We failed to recover *Zmadk1;2* double mutants, possibly due to lethality. However, plants homozygous for *Zmadk1;* and heterozygous for *Zmadk2* had larger shoot apical meristems even though the plants were dwarfed (Figures 4F and S12), indicating that they might directly affect meristem development. ADKs are predicted to function in cytokinin homeostasis to convert cytokinin nucleoside precursor (*N⁶*-(Δ²-isopentenyl) adenine riboside (iPR)) to the nucleotide (iPR 5’-phosphates (iPRPs)) forms (Schoor et al., 2011) (Figure 4H), and cytokinins control shoot meristem size (Giulini et al., 2004; Kurakawa et al., 2007). We therefore profiled hormones in immature ear primordia, and indeed found higher cytokinin nucleoside (iPR) and lower cytokinin nucleotide (iPRPs) levels in the mutants (Figures 4G and 4H). Meanwhile, the active cytokinin nucleobase forms (*cis*-zeatin (cZ) and *N⁶*-(Δ²-isopentenyl) adenine (iP)) were significantly upregulated in the mutants (Figures 4G and 4H). Active cytokinin promotes *WUS* expression and increases meristem size (Gordon et al., 2009). Taken together, our single-cell comparative analysis of stem cell over-proliferation mutants identified two subfamilies of carbohydrate kinase genes underlying either hypoxia signaling or cytokinin homeostasis (Figures 4E and 4K).

### Stem cell regulators identified from single-cell analysis were highly associated with yield traits

Maize stem cell activity is highly associated with ear yield traits (Lindsay et al., 2024). To test if our newly identified stem cell markers (Table S2) and genes differentially expressed in proliferating stem cells (Table S4) contribute to ear yield traits (Xu et al., 2021), we estimated the narrow-sense heritability (*h^2^*) of eight ear traits (Xu et al., 2021) and compared them to random distributions of *h^2^* generated from random gene subsets (Figure 5 A-D). We focused on markers within genes or 2kb upstream or downstream to identify potential associations between ear yield traits and gene function. We found that maize and arabidopsis conserved stem cell markers (Figures 2A and Table S2), explained significant *h^2^* for Cob Diameter and Cob Weight traits (Permutation P-value < 0.05; Figure 5A, Table S5)., and maize stem cell markers (Table S2) explained significant *h^2^* for Ear Length (Permutation P-value < 0.05; Figure 5B, Table S5). Furthermore, the differentially expressed genes (DEGs) in the proliferating mutant stem cells (Table S4) explained significant *h^2^* for Ear Row Number (Permutation P-value < 0.05; Figures 5C-D). These results were consistent for stem cells of both inflorescence meristems (Figure 5C) and spikelet pair meristems (Figure 5D), albeit the percentile of *h^2^* estimated from inflorescence meristems datasets (Table S4) was much higher (Figure 5C, Table S5).

**Figure 5:**
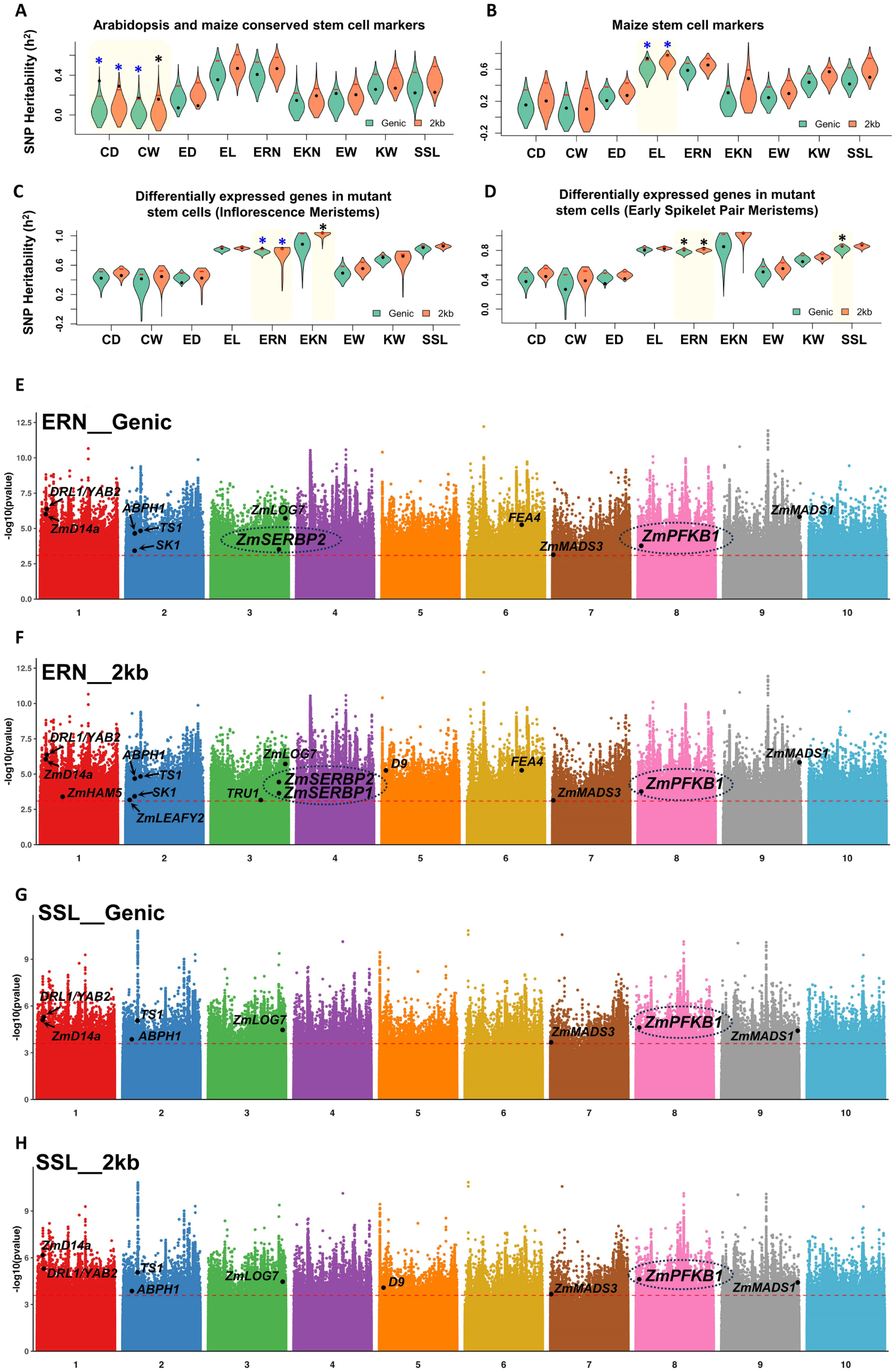
Stem cell regulators identified from single-cell analysis are highly associated with ear morphology trait. (A-D) Distributions of SNP heritability (*h^2^*) using genic (green violin plots) or 2kb upstream and downstream (orange violin plots) region; blue color * indicates *h^2^* values for stem cells marker genes (A-B) and DEGs in mutant cells (C-D) (black dots) for the given traits are greater than the top 5% permuted *h^2^* values (red bars) using 1,000 random subsets of maize genes. blue color * indicates *h^2^* values for stem cells marker genes (A-B) and DEGs in mutant cells (C-D) (black dots) for the given traits are greater than the top 10% permuted *h^2^* values using 1,000 random subsets of maize genes. *h^2^* values are reported in Table S5. (E-H) GWAS using SNPs in genic or 2kb upstream and downstream of genes reveals that newly identified stem cell regulators, *ZmSERBPs* and *ZmPFKB1* are highly associated with Ear Row Number (E-F) and/or Seed Set Length (G-H) traits. Red dash line indicates FDR threshold of 0.05. y axis indicates the –log10(p-value) (Table S5). Known maize developmental regulators passed FDR threshold of 0.05 were marked as a reference: *ZmLOG7*, *FEA4*, *ABPH1*, *DROOPING LEAF1*/*YABBY2* (*DRL1/YAB2*) (Strable et al., 2017), *ZmDWARF 14a* (*ZmD14a*) (Y. Liu et al., 2021), *TASSELSEED1* (*TS1*) (Acosta et al., 2009), *SILKLESS1* (*SK1*) (Hayward et al., 2016), *ZmMADS1*/*ZmMADS3* (Heuer et al., 2001), *ZmHAIRY MERISTEM5* (*ZmHAM5*) (Satterlee et al., 2020), *ZmLEAFY2* (Bomblies et al., 2003), and *TASSELS REPLACE UPPER EARS1* (*TRU1*) (Dong et al., 2017).

To further test if our newly identified stem cell markers (Table S2) and mutant stem cells DEG (Table S4) correspond to yield trait regulators, we performed GWAS analysis by comparing them against imputed whole genome sequencing (WGS) data (Grzybowski et al., 2023). This WGS data has higher coverage compared to the genotyping-by-sequencing data used in our previous study (Xu et al., 2021). From all genes with colocalized markers passing an FDR threshold of 0.05, we found that stem cell markers and DEGs in mutant stem cells were enriched for Ear Row Number, Ear Diameter, Ear Length, and Seed Set Length traits when considering both genic and +/- 2kb regions (p < 0.05, chi-square test; Figures 5E-F; Table S5). Additionally, we found significant enrichment for Ear Row Number when only considering markers in genic regions (p < 0.05, chi-square test; Table S5). Genes passing the FDR cutoff of 0.05 included *ZmSERBPs* (Figures 2D-F) and *ZmPFKB1* (Figure 3G-H, 4A-D) (Figures 5E-H; Table S5) newly identified stem cell regulators in this study, as well as known stem cell regulators *ZmLOG7* (Knauer et al., 2019), *FEA4* (Pautler et al., 2015), and *ABPH1* (Jackson and Hake, 1999). These findings suggest that candidate stem cell regulators identified via our single-cell analysis underly variation in maize yield traits.

## DISCUSSION

Our study overcame the challenge of recovering rare stem cells in single-cell profiling of maize and arabidopsis shoot tissues . Fine dissection of stem cell-enriched tissues, coupled with optimization of protoplast isolation, could be applied to capture rare and fragile shoot stem cells in other species. Our single-cell RNA-seq uncovered hundreds of uncharacterized marker genes that were highly co-expressed with *ZmCLE7* and *CLV3* in maize and arabidopsis, respectively. We validated their expression in stem cells using both traditional mRNA *in situ* hybridization and a spatial transcriptomics approach that can profile up to 100 genes at a time.

To ask if we could identify new stem cell regulators conserved across species, we compared our single-cell analysis between maize and arabidopsis. We identified conserved stem cell markers such as those encoding transcription factors and RNA-binding proteins. Many GO terms for stem cell markers were highly concordant between the two species, especially those related to gene expression regulation and metabolic processes. We functionally validated a family of *SERBP* RNA-binding protein-encoding genes, and found they control shoot meristem development in both maize and arabidopsis. Our GWAS analysis also found that maize *SERBPs* were associated with Ear Row Number, a yield associated trait. Future phenotypic studies of this trait could be conducted using our *Zmserbp* CRISPR mutants. SERBP1 and its orthologs are associated with ribosomes in human, drosophila, yeast, and cancer cells, and regulate protein translation and tumor progression (Anger et al., 2013; Ben-Shem et al., 2011; Brown et al., 2018; Martini et al., 2021; Muto et al., 2018). Coincidentally, ∼70% of the conserved stem cell markers were ribosomal-protein encoding genes. We validated the stem cell enriched expression of some of these genes, supporting the recent discovery of ribosome heterogeneity and specialization in stem cell differentiation (Barna et al., 2022; Genuth et al., 2022; Genuth and Barna, 2018). Future work will examine if SERBPs associate with ribosomes in plants, and may support a cross-kingdom conservation of their regulation of translation, as well as asking if ribosomal genes have specific biological roles in plant stem cells (Genuth et al., 2022).

We also profiled stem cell over-proliferation mutants, *fea3;Zmcle7* and *Bif3*, and identified candidate regulators that were differentially expressed in mutant stem cells. We functionally characterized families of redundant *PFKB* and *ADK* genes that are putative ZmWUS1 targets and have stem cell enriched expression. Analysis of *PFKB* expression suggests that expression of these genes responds to the hypoxic environment in stem cells. Hypoxia signaling is critical for pluripotency, proliferation, and differentiation of human stem cells and is of significant interest in stem cell therapeutics (Abdollahi et al., 2011; Di Mattia et al., 2021; Li et al., 2021). In tomato, *WUS* expression is repressed by *LITTLE ZIPPER* homolog *DEFECTIVE TOMATO MERISTEM* (*DTM*) (Xu et al., 2019), and *LITTLE ZIPPER* genes are regulated by hypoxia in arabidopsis (Weits et al., 2019). Since *ZmPFKBs* are candidate ZmWUS1 targets (Figure 4A), it would be interesting to test if the altered expression of *ZmPFKB* genes under hypoxic conditions might be due to direct regulation by a LITTLE ZIPPER-WUS transcriptional cascade in maize. Furthermore, supporting the potential importance of *PFKB* genes in development, our GWAS analysis found significant correlation for these genes with maize Ear Row Number and Seed Set Length yield component traits. In contrast to *ZmPFKBs*, the related *ZmADK* genes appear to function by a distinct mechanism by controlling cytokinin hormone homeostasis. Our results suggest that ZmWUS1 regulation of *ZmADK* genes controls active cytokinin levels to promote shoot meristem development.

Together, our studies establish a blueprint for identifying shoot stem cell regulators at single-cell resolution to inform functional analyses (Xu et al., 2022). By providing functional validation and ear yield trait association analysis, we identified new regulators of shoot stem cells that could be exploited to modify plant architecture and improve crop productivity.

## ACKNOWLEDGMENTS

We thank Samik Bhattacharya, Jasper Kläver, and Nachiket Kashikar from Resolve Biosciences for spatial transcriptomic service, Margaret Woodhouse and John Portwood for hosting our single-cell datasets on MaizeGDB, Drs. Michael Scanlon and James Satterlee for advice on shoot protoplasting methods, Liang Dong and Dr. Fang Yang for sharing ZmWUS1 tsCUT&Tag peak files (Dong et al., 2022), Dr. Munenori Kitagawa for sharing *ap1;cal* double mutant seeds, Dr. Karen Koch for inspiration in testing the hypoxia hypothesis, Dr. Andrea Schorn for sharing hypoxia chambers. We thank support from the Cold Spring Harbor Laboratory farm management team, single-cell core, and NGS facility. We acknowledge funding support from NSF (IOS-1833182, IOS-1445025, and IOS-1934388). X.X. acknowledges support from California Agricultural Experiment Station (AES). J.G. acknowledges support from R01 LM012736 and R01 MH113005. A.G. acknowledges support from NSF (IOS 2026561); M.P. acknowledges support from the William Randolph Hearst Scholarship. B.H. acknowledges support from The Robertson Research Fund, CSHL School for Biological Sciences.

## AUTHOR CONTRIBUTIONS

X.X. performed all experimental procedures and data analysis, except for those listed below, and wrote the draft of the manuscript. M.P. performed the maize and arabidopsis co-expresslog analysis and differential expression analysis between wild type and stem cell mutants. B.R. performed narrow-sense heritability and GWAS analyses. E.D.A. made and characterized *fea3*;*Zmcle7* double mutants. M.K. and Y.T. performed the hormone measurement under the supervision of H.S. B.H. assisted with the Slingshot trajectory analysis. A.G. generated *Bif3* mutant seeds. D.J. and J.G. supervised the research. D.J. co-wrote the manuscript, and all authors edited it.

## STAR**↔**METHODS

### RESOURCE AVAILABILITY

#### Lead Contact

Further information and requests for resources and reagents should be directed to and will be fulfilled by the Lead Contact, David Jackson.

### Materials Availability

Requests for materials should be directed to Lead Contact, David Jackson.

### Data and Code Availability

All raw high-throughput sequencing data have been deposited in NCBI’s Sequence Read Archive (SRA). SRA IDs are listed in Table S1. This study used codes from published software described in QUANTIFICATION AND STATISTICAL ANALYSIS.

## EXPERIMENTAL MODEL AND SUBJECT DETAILS

Maize plants were grown in standard field or greenhouse conditions (Je et al., 2016). Reference B73 inbred plants were used for wild type ear tip single-cell experiments (four biological replicates), mRNA *in situ* hybridization, spatial transcriptomic analysis, and gene expression analysis of hypoxia response. *fea3* (B73 background) (Je et al., 2016) was crossed to *Zmcle7* (B73 background) (Rodriguez-Leal et al., 2019) to generate double mutants for single-cell profiling (three biological replicates). *Bif3* (A619 background) (Chen et al., 2021) homozygous plants used for single-cell RNA-seq (two biological replicates) Genotyping *fea3*, *Zmcle7*, and *Bif3* was as previously described (Chen et al., 2021; Je et al., 2016; Rodriguez-Leal et al., 2019). *Atserbp* mutants were obtained from ABRC. Genotyping primers were listed in Table S6. arabidopsis *ap1;cal* (Col) double mutants were grown in long day conditions in a growth chamber. The inflorescence apices were harvested for protoplasting.

A multiplex CRISPR/Cas9 strategy was used to knockout *ZmSERBP*, *ZmPFKB*, and *ZmADK* genes following *Agrobacterium*-mediated transformation of B104 embryos. Guide RNAs (sgRNAs) were designed based on B73 V3 or V4 reference genome using CRISPR-P (http://crispr.hzau.edu.cn/CRISPR2/), and their specificity in B104 confirmed (MaizeGDB) by blast. Sequences of sgRNAs were shown in Figure S10. The sgRNAs arrays were synthesized by Gene Universal Inc. with *Hind*III sites at the ends for cloning into pCPB-ZmUbi-hspCas9 vector (Liu et al., 2020). CRISPR constructs were transformed into *Agrobacterium* (EHA105) for transformation at the Iowa state transformation facility. Transformation events were obtained and analyzed by PCR amplicon barcode genotyping (L. Liu et al., 2021a). Barcode PCR primers were listed in Table S6.

## METHOD DETAILS

### Protoplast preparation and 10x Genomics library construction and sequencing

For wild type B73 and *fea3;Zmcle7* ear tip single-cell RNA-seq, the ear tips including inflorescence and early spikelet pair meristems were finely dissected. Two wild type replicates and one replicate of *fea3;Zmcle7* were generated using a MES buffer based enzyme mix as previously reported (Ortiz-Ramírez et al., 2018; Xu et al., 2021). Protoplasts were purified using OptiPrep Gradient Medium by modifying a protocol described in Application Sheet C18; 6th edition, January 2018 (protocol will be provided upon request). In detail, (1) We skipped solution A and steps 2b-1 and 2b-2. (2) In step 2b-3, we replaced solution B with MES buffer based enzyme mix and isolated protoplasts as previously reported up to the point of FACS purification (Ortiz-Ramírez et al., 2018; Xu et al., 2021). (3) In step 2b-4, we replaced solution C with our MES based wash buffer as previously reported (Ortiz-Ramírez et al., 2018; Xu et al., 2021). (4) Protoplasts were harvested from the top layer into a 15ml falcon tube. (5) Wash buffered was gently added up to 6ml total volume, and the tubes were centrifuged for 2min at 500g at 4^0^C. (6) The supernatant was then removed to keep ∼ 30-200ul leftover depending on protoplast pellet size. (7) Protoplasts were then gently suspended using a P200 pipette with the pipette tip cut off using a razor blade. (8) Aliquots of protoplast were immediately stained by trypan blue and counted under a DIC microscope. Another two replicates of B73 and *fea3;Zmcle7*, and all replicates of *ap1;cal* and *Bif3* were generated using a MOPS buffer based enzyme mix and washing buffer as reported in (Satterlee et al., 2020) with cell isolation steps optimized as listed below: after tissue digestion with MOPS buffer based enzyme mix, instead of using OptiPrep Gradient Medium purification, we first filtered the digested protoplasts using a 40μm pluriStrainer (pluriSelect, Catalog No. 43-50040-50) sitting on a 100x15mm petri dish (VWR, Catalog No. 25384-342), rather than 50ml falcon tube to reduce the falling distance after protoplasts were filtered. The filtered protoplasts were then gently poured into an ice-cold 15ml falcon tube to avoid shearing force that could damage fragile stem cells. The protoplast suspension was then centrifuged for 2 min at 300g at 4^0^C using a swing bucket centrifuge to collect cell pellets. The washing steps were also performed at 4^0^C. The enzyme mix supernatant was then carefully removed by pipetting without disturbing the cell pellets. Next, 1ml ice cold MOPS based washing buffer was carefully added onto the protoplast pellets and the pellets were gently resuspended using a 1-ml pipette with the tip cut off using a razor blade. An additional 1ml ice cold MOPS based washing buffer was added, and again was gently mixed using a 1ml pipette tip cut off by a razor blade. Finally, 8ml ice cold MOPS based washing buffer was added and the tube was gently inverted 3 times to wash the protoplasts. The protoplasts were then gently poured into a 30μm pluriStrainer (pluriSelect, Catalog No. 43-50030-50) sitting on a 100x15mm petri dish (VWR, Catalog No. 25384-342). The filtered protoplasts were carefully poured into a new ice-cold 15ml falcon tube. The tube was then centrifuged for 2 min at 300g. The supernatant was then carefully removed by pipetting, keeping ∼ 30-200μl leftover depending on protoplast pellet size. Cell pellets were not disturbed during the removal of washing buffer. The protoplasts were then gently suspended using a P200 pipette with the pipette tip cut off using a razor blade. Protoplast aliquots were immediately stained with trypan blue and counted under a DIC microscope. High viability (≥ 70%) protoplasts based on trypan blue staining were immediately loaded into a 10x Genomics Chromium System using V3 chemistry kits. Single-cell RNA-seq libraries were sequenced by Illumina PE150 reads with ∼ 400M paired-end reads per library. We successfully recovered stem cells using both methods, thus we used all data for downstream analysis. Raw sequencing data were deposited in NCBI’s Sequence Read Archive (SRA). SRA IDs are listed in Table S1.

### Scanning electron and Confocal microscopy

Scanning electron microscopy was performed on fresh tissues of maize and arabidopsis using a JEOL Benchtop SEM JCM-7000 under high vacuum settings. *pZmWUS1*-mRFP lines (Je et al., 2016) were imaged using a ZEISS LSM 780 confocal microscope. The DAPI channel was used for capturing autofluorescence.

### mRNA *in situ* hybridization and spatial transcriptomic analysis

mRNA *in situs* and probe synthesis were conducted as previously described (Jackson et al., 1994; Xu et al., 2021). Primer sequences for all genes are listed in Table S6.

Spatial transcriptomics was performed using Resolve Biosciences (Germany) platform as previously described (Guillotin et al., 2023) except using wild type B73 paraffin-embedded ear inflorescence tissues.

### Gene expression analysis of hypoxia response

Wild type B73 ear primordia (2-4 mm) were dissected and put on shoot culture medium (Irish and Karlen, 1998) in petri dishes sealed with 3M micropore tape (REF 1530-0) to allow air exchange. The petri dishes were then incubated in either low oxygen (2%) or normal oxygen (21%) incubators (Thermo Scientific, Heracell™ 150i) at 28^0^C for 24hrs. Tissues were then immediately frozen in liquid nitrogen and RNA extracted using a Zymo Research Direct-zol RNA MiniPrep kit followed by cDNA synthesis (Invitrogen SuperScript III). qPCR primers were designed by IDT PrimerQuest Tool and qPCR assays were performed with a QuantStudio™ 5 Real-Time PCR System. qPCR primers were listed in Table S6.

### Cytokinin quantification

Cytokinins were extracted and semi-purified from maize immature ear primordia as described (Kojima et al., 2009). They were quantified using an ultra-performance liquid chromatography (UPLC)-tandem quadrupole mass spectrometer (ACQUITY UPLC System/Xevo-TQS; Waters Corp.) with an octadecylsilyl (ODS) column (ACQUITY UPLC HSS T3, 1.8 µm, 2.1 mm × 100 mm, Waters Corp.).

### Meristem size measurement

Meristem size measurements were performed as previously described (Wu et al., 2020). In brief, shoot apical meristems from maize or arabidopsis seedlings were dissected and fixed in FAA (10% formalin, 5% acetic acid, and 45% ethanol) overnight at 4^0^C. The fixed tissues were then dehydrated in 70%, 85%, 95%, 100%, and 100% ethanol for 60 min each at room temperature.

Then the dehydrated tissues were immersed in a 1:1 ethanol-methyl salicylate solution for an additional 60 min. The tissues were then cleared with methyl salicylate (three times 60 min with fresh methyl salicylate). The cleared shoot apical meristems were imaged with a Nikon DS-Ri2 DIC microscope.

## QUANTIFICATION AND STATISTICAL ANALYSIS

### Single-cell RNA-seq analysis, clustering, and selection of marker genes

Sequencing reads of all maize single-cell RNA-seq samples were first aligned to an updated maize v3 reference genome (Xu et al., 2021) using Cell Ranger3.1.0 (10X Genomics). Downstream cluster processing was performed in R using the Seurat V4.0.3 package (Hao et al., 2021). Genes expressed in less than 3 cells were removed. For maize wild type and mutant ear tip datasets, we kept cells with 1,000 to 15,000 expressed genes and 5,000 to 150,000 unique molecular identifiers (UMIs). Then reads of individual samples were log-normalized and 2000 highly variable genes were used for downstream analysis. Replicates were integrated using Seurat default *FindIntegrationAnchors* followed by *IntegrateData* functions. The integrated matrix was then scaled using the *ScaleData* function. Principal Component analysis (PCA) was then performed by *RunPCA* function. The top principal components were selected based on Elbow plot-based quantitative approach (https://hbctraining.github.io/scRNA-seq/lessons/elbow_plot_metric.html). Clusters were then identified with *FindNeighbors* and *FindClusters* functions. To generate subclusters for cluster 6 in wild type B73 ear tip and cluster 2 in integrated datasets of wild type B73 and *Bif3*, the Seurat *FindSubCluster* function was used with a resolution of 0.2. Three cell cycle G2/M clusters (7, 9, and 13) were merged into one G2/M cluster. Cluster-specific markers were identified using the Seurat *FindAllMarkers* function under parameters of p_val_adj < 0.01 and logfc.threshold = 0.5 (Tables S1).

For arabidopsis *ap1;cal* inflorescence apex datasets, the same analysis workflow was applied to generate clusters and identify cluster-specific markers, except sequencing reads were aligned to the arabidopsis TAIR10 reference genome, and cells with 1,000 to 100,000 UMIs and 500 to 12,000 expressed genes were kept.

### Single-cell RNA-seq cross-species co-expression analysis

Single-cell co-expression analysis was performed in R. Briefly, the gene count matrix of all the cells from the Seurat object of either maize wild type ear apex or arabidopsis *ap1;cal* was extracted. Then the Pearson correlation coefficient value between genes of interest (such as *ZmCLE7* in maize or *CLV3* in arabidopsis) and all other expressed genes was then calculated using *cor* function (Table S2)(Yang et al., 2021). The top 300 highly co-expressed genes were identified in both maize and arabidopsis by ranking the Pearson correlation coefficient values (Table S2).

We performed a BLAST search of the top 300 maize genes that were highly co-expressed with *ZmCLE7* in Phytozome to identify the top 10 homologs of each gene based on E-value in arabidopsis. Similarly, the top maize homologs of the top 300 *CLV3* highly co-expressed genes in arabidopsis were identified. The overlap between maize and arabidopsis genes was then reciprocally checked, and a chi-squared test was performed to determine the significance of overlaps. The overlapped genes were concluded as highly enriched stem cell markers between maize and arabidopsis (Table S2).

### Comparison of arabidopsis single-cell RNA-seq and FACS-microarray data

Single-cell co-expression analysis for *CLV3* and *FIL* was performed in R as mentioned above. Top 300 co-expressed markers were selected to check the overlap with domain-specific genes identified in Yadav et al., 2019 using CLV3 (genes reported in S5) and FIL (genes reported in S7). A chi-squared test was performed to determine the significance of overlaps (Table S1).

### Single-cell RNA-seq cross-species cell type comparison

To compare the single-cell data of maize and arabidopsis at cell type levels, we directly integrated all the cells from maize wild type ear tip and arabidopsis *ap1;cal* datasets. Cross-species cell integration requires sufficient high-confidence functionally orthologous gene pairs. To improve the integration by expanding the shared gene space, we used the co-expression proxies identified by EPIPHITES (Passalacqua and Gillis, 2023) to identify genes pairs between maize and arabidopsis (https://gillislab.shinyapps.io/epiphites/). Briefly, EPIPHITES identifies pairs of genes across two species that share an orthology relationship and are highly similar in expression profile. This approach identified 6784 gene pairs. We converted the maize V4 IDs to V3 IDs and 6387 gene pairs were preserved.

Next, we used the 6387 gene pairs to perform Seurat integration with a similar workflow as mentioned above. The cell clusters were annotated either using known markers or projecting back to the original wild type ear tip or arabidopsis *ap1;cal* clusters (Figure S4B-C). Conserved cell types were identified based on the comparable cells between maize and arabidopsis. Cluster-specific conserved markers were identified using Seurat *FindConservedMarkers* function under parameters of p_val_adj < 0.01 and logfc.threshold = 0.25 (Tables S3) (Hao et al., 2021).

### Single-cell RNA-seq comparison between wild type and stem cell mutants

We first integrated the wild type ear tip datasets with either *fea3;Zmcle7* or *Bif3* using Seurat integration (Stuart et al., 2019) pipeline. The genotype information was added to the metadata of the integrated object to separate the wild type and mutant samples. After integration, clusters were annotated using known markers. We focused on the central zone stem cell clusters to perform differential expression analysis between wild type and mutants using the Seurat FindMarkers functions default parameters with ident.1 = “wild type”, ident.2 = “*fea3;Zmcle7*” or “*Bif3*”. To identify high confidence candidate differentially expressed genes across both mutants, we used Fisher’s method to combine the uncorrected *p*-values from both differential gene analyses. We then performed multiple hypothesis correction using the Bonferroni correction at an alpha of .05 for all genes. This correction is highly stringent, and post-correction results in a 5% chance of a single false positive across all genes.

### Gene Ontology Analysis

Gene Ontology (GO) analysis was performed using http://geneontology.org. The enrichment in Biological Process, Molecular Function, and Cellular Component was checked individually. Over-represented GO terms with FDR P<0.05 were analyzed and reported in Tables S2 and S3.

### Narrow-sense heritability

For additive genetic variance estimation, we estimated narrow-sense heritability (*h^2^*) from the marker subsets (Speed et al., 2012). To assess whether the derived heritability for a specific trait significantly exceeded chance, we conducted 1000 permutations using random subsets of maize genes and estimated heritability accordingly. In each permutation, genes containing at least one marker within their genic region equal to the given set were randomly chosen. A target set was deemed significant for a particular trait if its observed heritability surpassed the upper 5th percentile threshold of permuted values.

### GWAS

We investigated significant colocalization patterns within GWAS data pertaining to maize ear traits. Utilizing the best linear unbiased predictors (BLUPs) derived from nine distinct ear phenotypes, as outlined in (Rice et al., 2020), we conducted our analysis. We employed publicly available imputed whole genome sequencing data (Grzybowski et al., 2023). A generalized linear model incorporating the first three principal components derived from a randomly selected subset of 100,000 SNP markers was executed genome-wide. GWAS was conducted across the entire marker spectrum, encompassing both SNP and short INDEL markers. To assess colocalization significance, a chi-square test was employed to ascertain whether the number of a-priori genes deemed significant was greater than the total count of genome-wide significant genes. Significant genes were determined if they had markers surpassing an FDR-adjusted threshold of 0.05 within the genic region or 2kb upstream and downstream.

---

**Figure S1:**
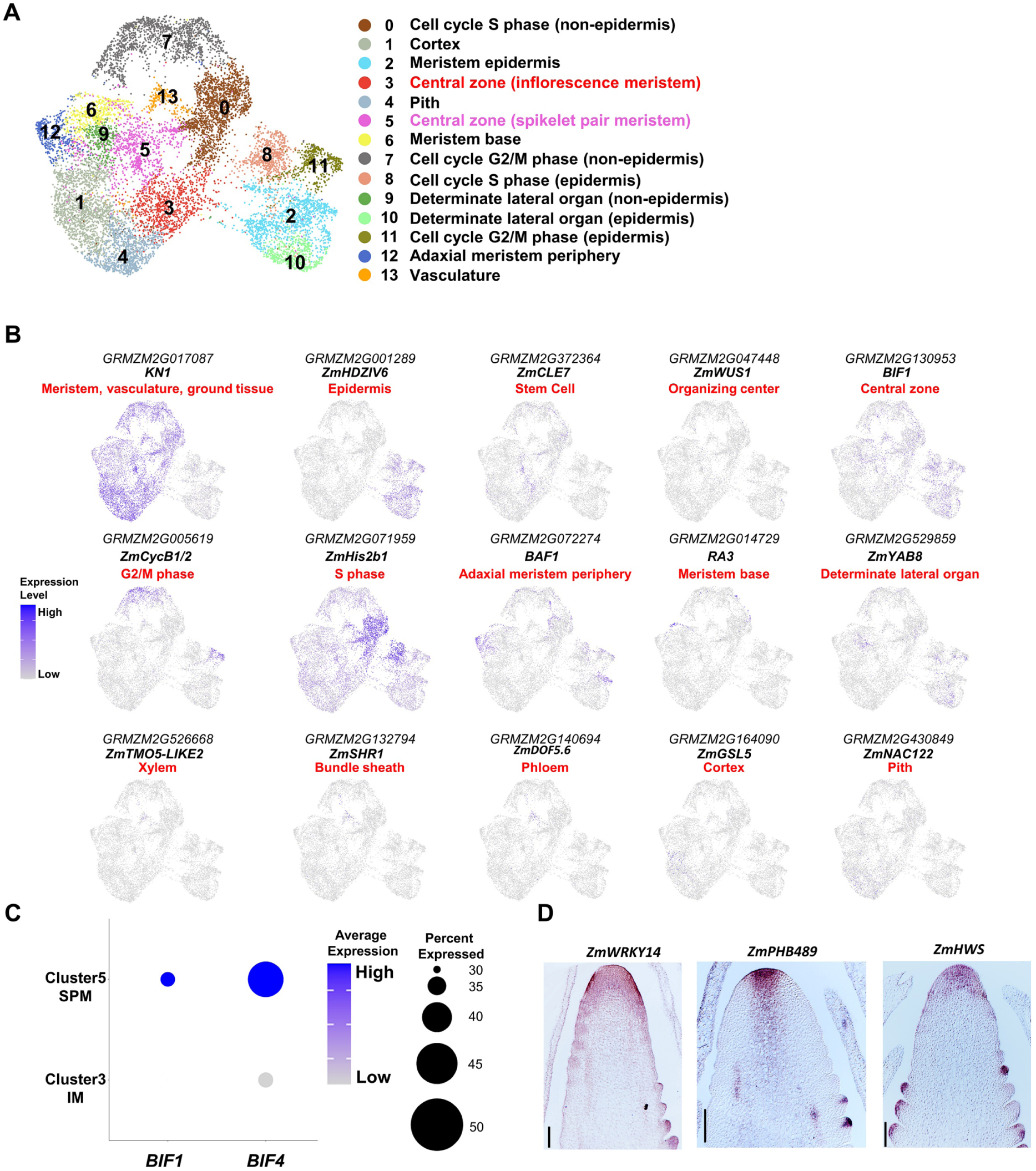
Annotation and validation of wild type ear tip cell clusters. (A) 14 distinct single-cell RNA-seq clusters of maize wild type ear tip. (B) UMAP plots of marker genes predicting cluster identities, *KN1*, meristem (clusters 3, 5, 6, and 12), ground tissue (clusters 1 and 4) and vasculature (cluster 13); *ZmHDZIV6*, epidermis (clusters 2 and 10); *ZmCLE7*, stem cells (clusters 3 and 5); *ZmWUS1*, organizing center (clusters 3 and 5); *BIF1*, spikelet pair meristem (cluster 5); *ZmCYCLINB1;2* (*ZmCYCB1;2*), G2/M phase (clusters 7 and 11); *ZmHISTONE2b1* (*ZmHis2b1*), S phase (clusters 0, 8); *BARREN STALK FASTIGIATE1* (*BAF1*), adaxial meristem periphery (cluster 12); *RA3*, meristem base (cluster 6); *ZmYABBY8* (*ZmYAB8*), determinate lateral organ (cluster 9); *ZmTARGET OF MONOPTEROS5-LIKE2* (*ZmTMO5-LIKE2*), xylem (cluster 14); *ZmSHR1*, bundle sheath (cluster 14); maize Dof-type zinc finger DNA-binding family protein (*ZmDOF5.6*), phloem (cluster 14); homolog of the Gibberellic Acid Stimulated Transcript-like (GAST-like) gene (*ZmGSL5*) (Zimmermann et al., 2010), cortex (cluster 1); *ZmNO APICAL MERISTEM DOMAIN CONTAINING* (*NAC*) *TRANSCRIPTION FACTOR 122* (*ZmNAC122*), pith (cluster 4). Color scale indicating normalized expression level. (C) Dot plots showing enriched expression of axillary meristem marker genes *BIF1* and *BIF4* in cluster 5 (thus annotated as spikelet pair meristem), rather than cluster 3 (thus annotated as inflorescence meristem). (D) mRNA *in situ* hybridization of *ZmCLE7* highly co-expressed markers showed enriched expression in the stem cells. Scale bar = 100µm.

**Figure S2:**
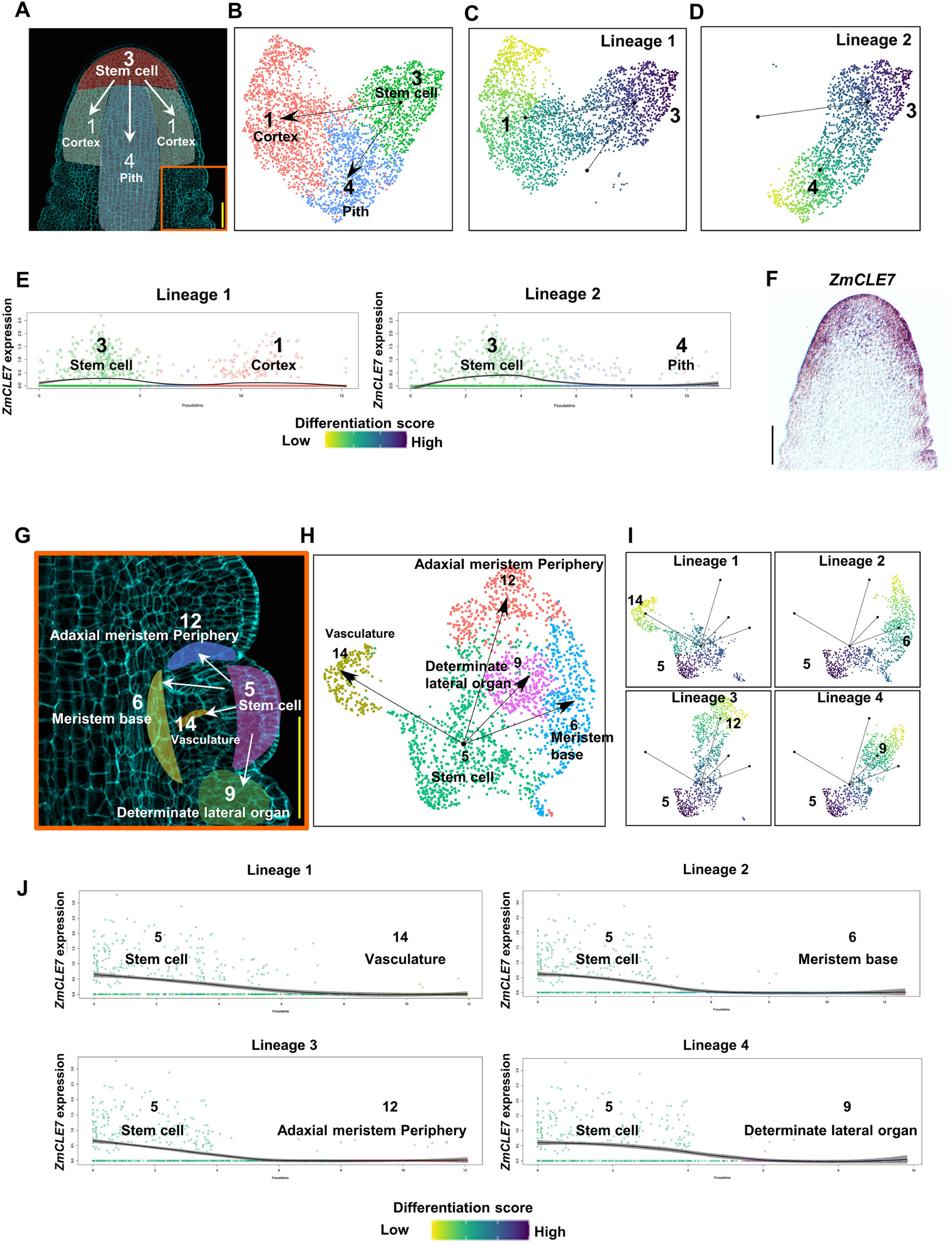
Developmental trajectory of wild type ear tip. (A-D) wild type ear inflorescence meristem (A) has two branching trajectories (B) from undifferentiated central zone stem cells to either cortex (C) or pith (D). (E) *ZmCLE7* showing dynamic expression along the two trajectory lineages in C (left panel) and D (right panel). (F) mRNA *in situ* hybridization of *ZmCLE7*. (G-J), wild type ear spikelet pair meristem (G) has two four branching trajectories (H-I) from undifferentiated central zone stem cells to other differentiated cell types. (J) *ZmCLE7* showing dynamic expression along the four branching trajectory lineages in H. Scale bar = 100µm in (A), (F), and (G). The region within orange box in (A) is shown in (G).

**Figure S3:**
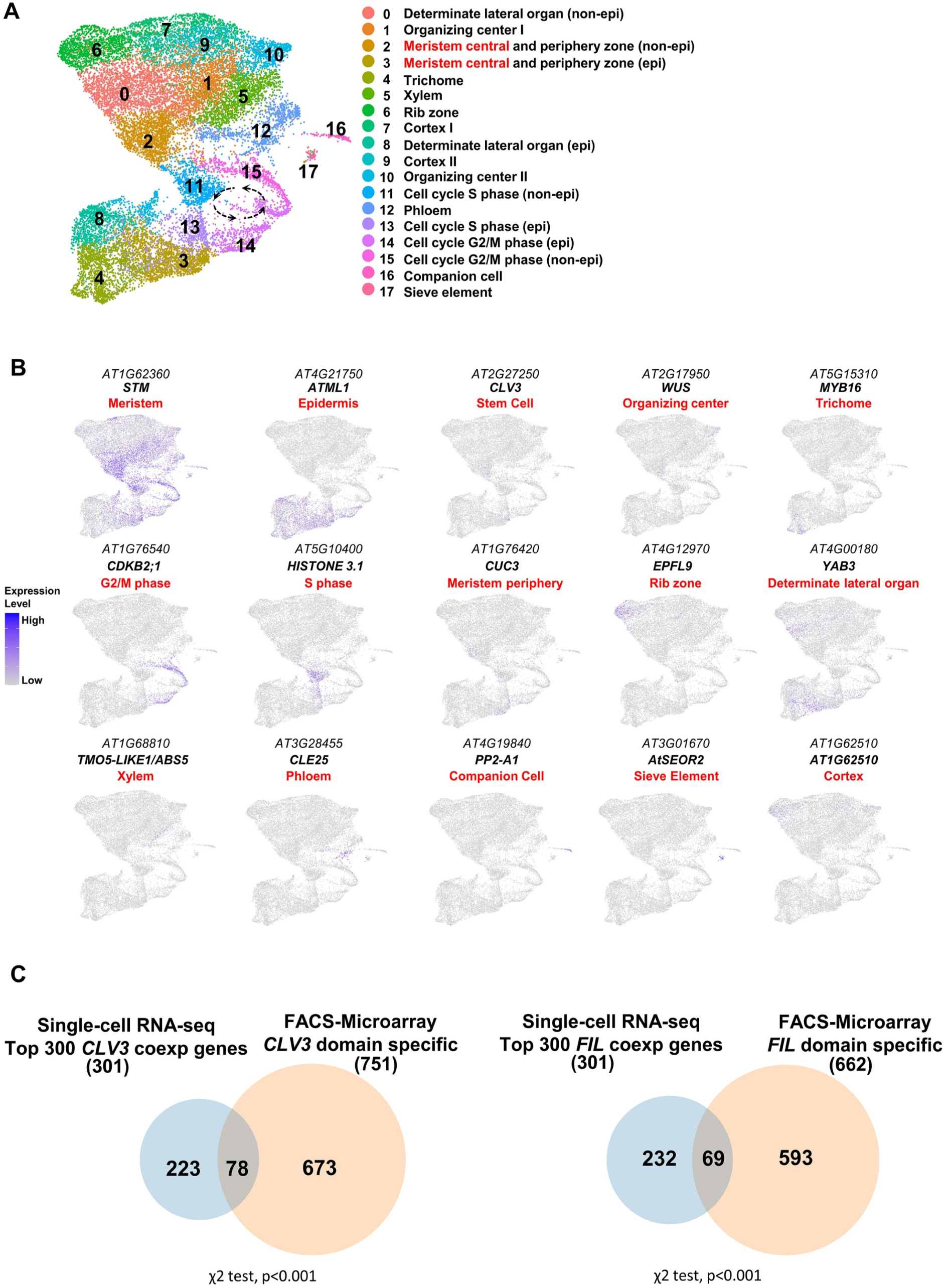
Annotation of arabidopsis shoot tip single-cell clusters. (A) 18 distinct single-cell RNA-seq clusters of arabidopsis *ap1;cal* mutant. (B) UMAP plots of marker genes predicting cluster identities, *STM*, meristem (clusters 3, 2, 1, 10), ground tissue (clusters 7 and 9) and vasculature (clusters 5, 12, 16, and 17); *ATML1*, epidermis (clusters 3, 4 and 8); *CLV3*, stem cells (part of clusters 3 and 2); *WUS*, organizing center (clusters 1 and part of clusters 2 and 10); *MIXTA-like* Transcription factor *MYB16* (*MYB16*)(Oshima and Mitsuda, 2013), Trichome (cluster 4); *CYCLIN-DEPENDENT KINASE B2;1* (*CDKB2;1*), G2/M phase (clusters 14 and 15); *HISTONE 3.1*, S phase (clusters 11, 13); *CUP SHAPED COTYLEDON3* (*CUC3*), meristem periphery (part of clusters 3 and 2); *EPIDERMAL PATTERNING FACTOR-like protein 9* (*EPFL9*), rib zone (cluster 6); *YABBY3* (*YAB3*), determinate lateral organ (clusters 0 and 8); *TMO5-LIKE1/ABS5*, xylem (cluster 5); *CLAVATA 3 (CLV3)/EMBRYO SURROUNDING REGION 25* (*CLE25*), phloem (cluster 12); *PHLOEM PROTEIN 2-A1* (*PP2-A1*), companion cell (cluster 16); *Arabidopsis THALIANA SIEVE ELEMENT OCCLUSION-RELATED 2* (*AtSEOR2*), sieve element (cluster 17); *AT1G62510*, cortex (clusters 7 and 9). (C) Venn diagram showing significant overlap between single-cell RNA-seq and FACS-microarray date for *CLV3* and *FIL*.

**Figure S4:**
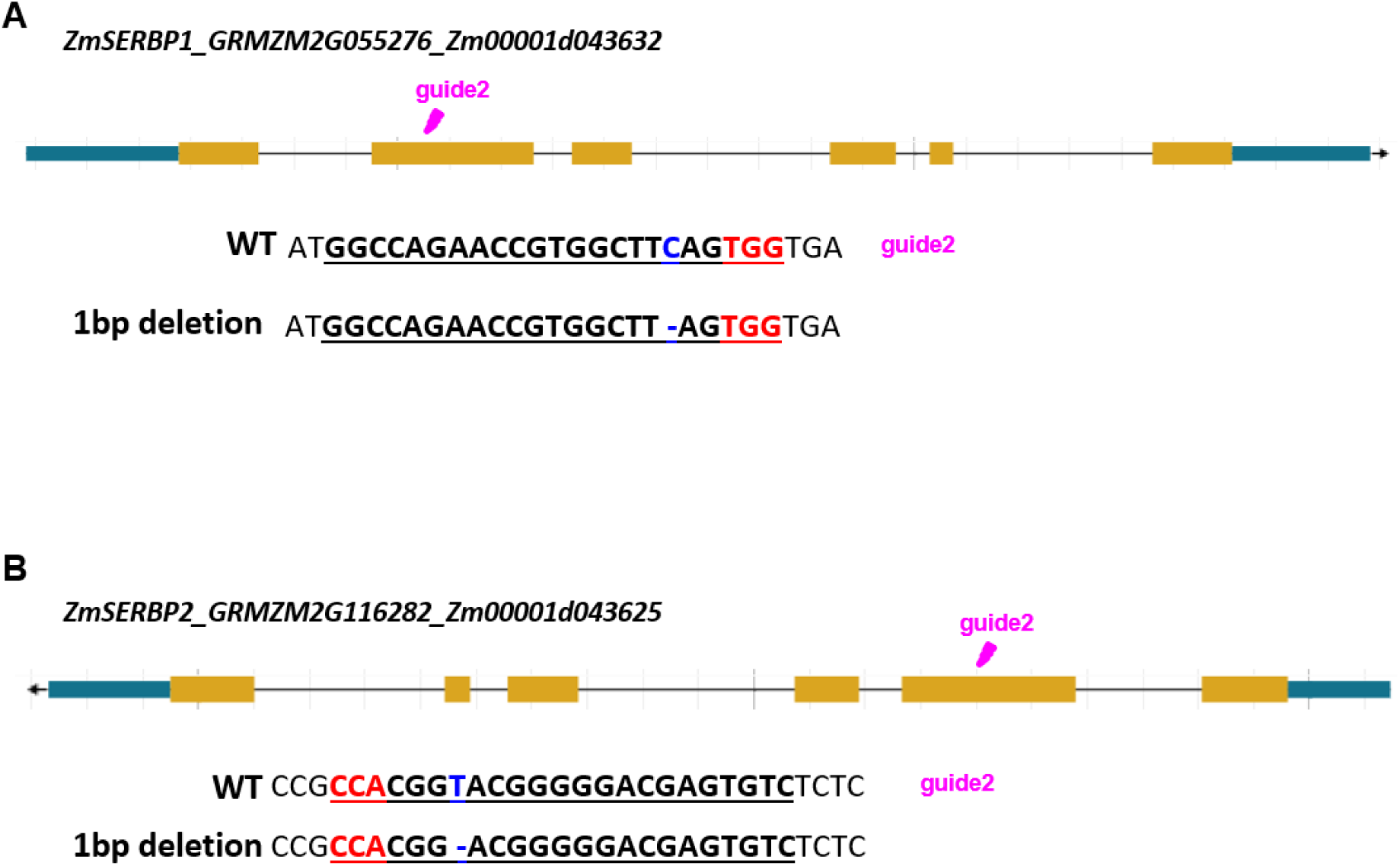
sgRNA and edits for *ZmSERBPs*. CRISPR gRNA-sequences and edits for *ZmSERBP1* (A) and *ZmSERBP2* (B)

**Figure S5:**
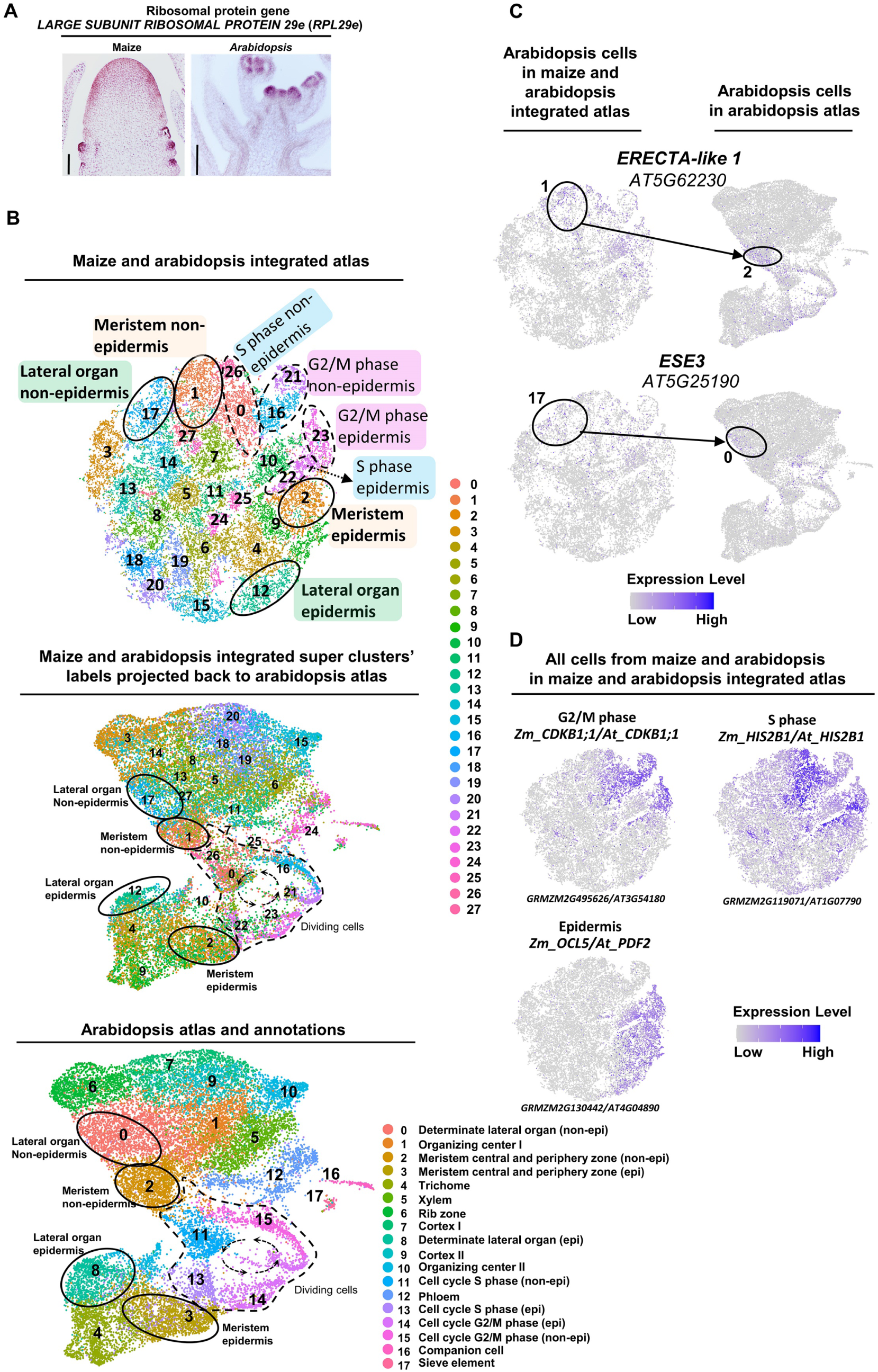
Cross-species analysis of arabidopsis *ap1;cal* and maize wild type B73 single-cell datasets. (A) mRNA *in situ* hybridization of conserved stem cell marker gene, *RIBOSOMAL PROTEIN L29e* (*RPL29e*), in both maize and arabidopsis. (B) Annotating maize-arabidopsis conserved super-clusters with arabidopsis cluster annotations (meristem, clusters 1 and 2; determinate lateral organ, clusters 12 and 17; G2/M phase, clusters 16, 21, and 23; and S phase, clusters 0, 22, and 26). (C) *t*-SNE plot of maize-Arabidopsis conserved marker genes in meristem (*ERECTA-like1*) and determinate lateral organ (*ETHYLENE AND SALT INDUCIBLE 3* (*ESE3*)) clusters. Color scale indicating normalized expression level. (D) *t*-SNE plots of epidermis and dividing cell marker genes. Color scales indicate normalized expression level.

**Figure S6:**
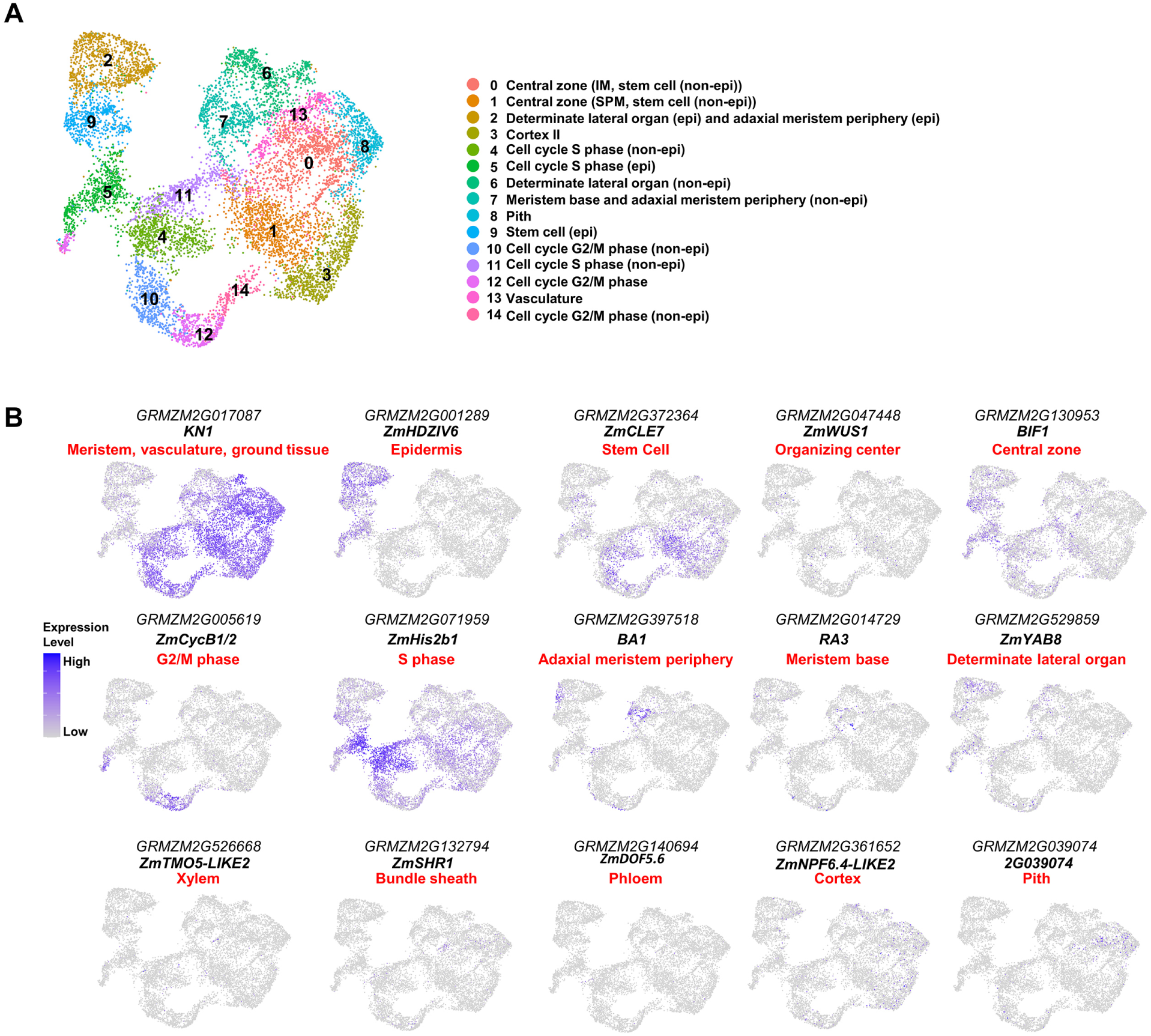
Annotation of *fea3;Zmcle7* ear tip cell clusters. (A) 15 distinct clusters of maize *fea3;Zmcle7* ear tip. (B) UMAP plots of marker genes predicting cluster identities, *KN1*, meristem (clusters 0, 1, and 7), ground tissue (clusters 3 and 8) and vasculature (cluster 13); *ZmHDZIV6*, epidermis (clusters 2 and 9); *ZmCLE7*, stem cells (clusters 0, 1 and 9); *ZmWUS1*, organizing center (clusters 0, 1 and 9); *BIF1*, spikelet pair meristem (cluster 1 and 9); *ZmCycB1/2*, G2/M phase (clusters 10, 12 and 14); *ZmHis2b1*, S phase (clusters 4 and 11); *BARREN STALK1* (*BA1*), adaxial meristem periphery (part of cluster 7 and 2); *RA3*, meristem base (part of cluster 7); *ZmYAB8*, determinate lateral organ (cluster 6 and part of cluster 2); *ZmTMO5-LIKE2*, xylem (cluster 13); *ZmSHR1*, bundle sheath (cluster 13); *ZmDOF5.6*, phloem (cluster 13); *ZmNITRATE TRANSPORTER 1/PEPTIDE TRANSPORTER FAMILY 6.4-LIKE 2* (*ZmNPF6.4-LIKE2*), cortex (cluster 3); *2G039074*, pith (cluster 8). Color scale indicating normalized expression level.

**Figure S7:**
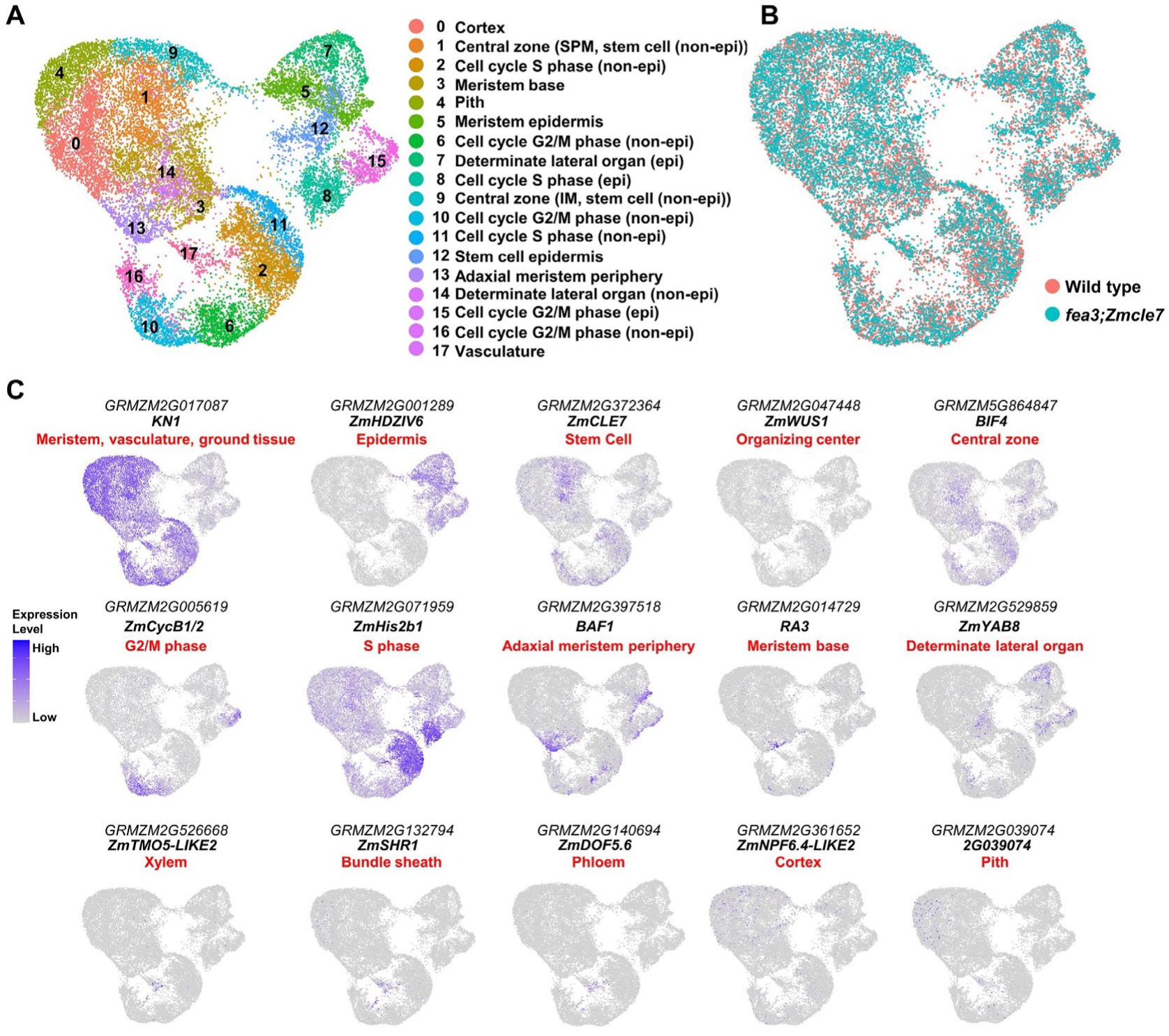
Integration of wild type and *fea3;Zmcle7* ear tip cell clusters. (A-B) UMAP plots showing 18 cell clusters (A) after integrating maize wild type and *fea3;Zmcle7* ear tip cells (B). (C) UMAP plots of marker genes predicting cluster identities, *KN1*, meristem (clusters 1, 9, 3, and 13), ground tissue (clusters 0 and 4) and vasculature (cluster 17); *ZmHDZIV6*, epidermis (clusters 5, 7 and 12); *ZmCLE7*, stem cells (clusters 1, 9, and 12); *ZmWUS1*, organizing center (clusters 1, 9, and 12); *BIF4*, spikelet pair meristem (cluster 1 and 12); *ZmCycB1/2*, G2/M phase (clusters 6, 10, 15 and 16); *ZmHis2b1*, S phase (clusters 2, 8 and 11); *BAF1*, adaxial meristem periphery (clusters 5 and 13); *RA3*, meristem base (part of cluster 3); *ZmYAB8*, determinate lateral organ (clusters 7 and 14); *ZmTMO5-LIKE2*, xylem (cluster 17); *ZmSHR1*, bundle sheath (cluster 17); *ZmDOF5.6*, phloem (cluster 17); *ZmNPF6.4-LIKE2*, cortex (cluster 4); *2G039074*, pith (cluster08). Color scale indicating normalized expression level.

**Figure S8:**
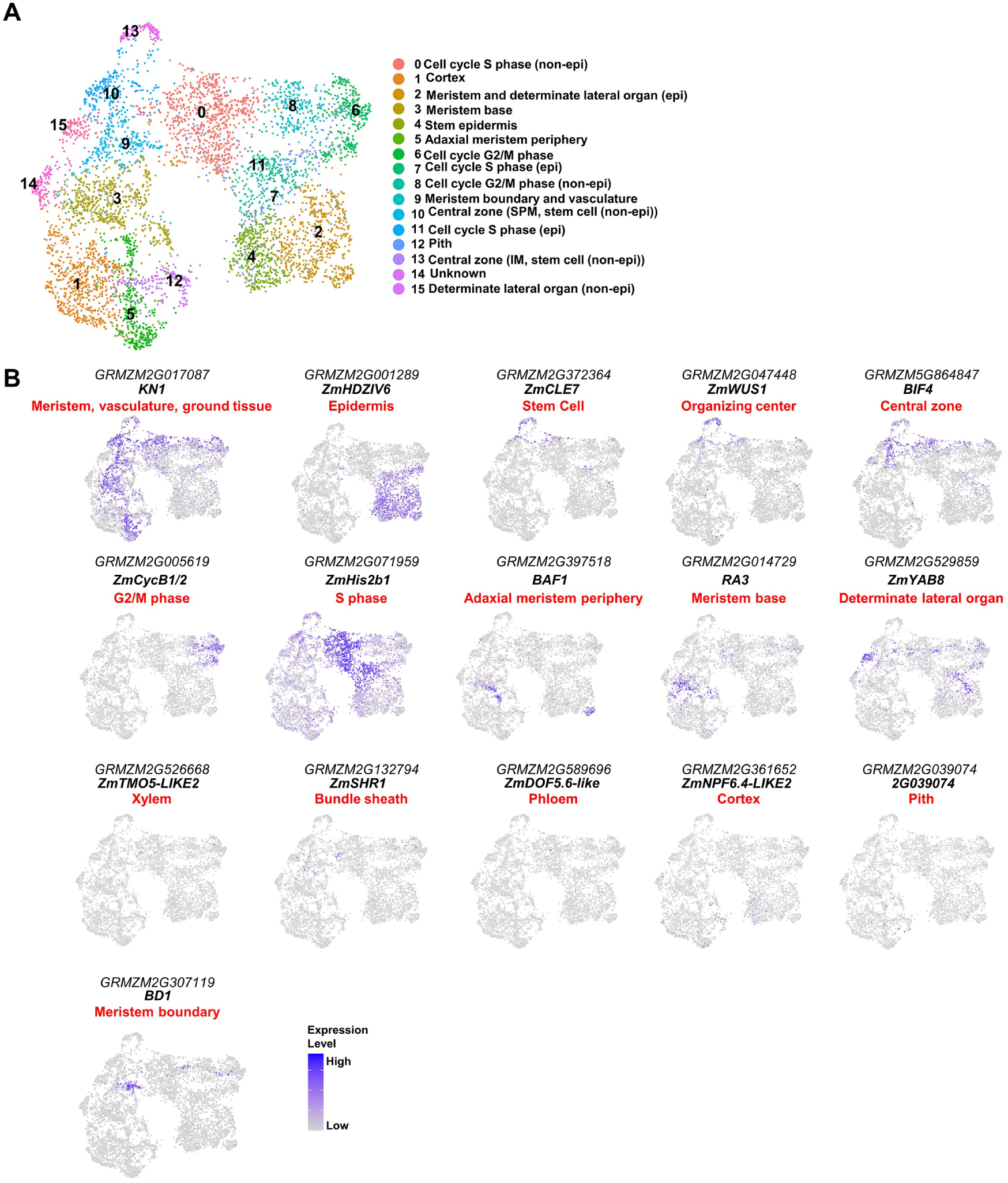
Annotation of *Bif3* ear tip cell clusters. (A) 16 distinct clusters of *Bif3* ear tip. (B) UMAP plots of marker genes predicting cluster identities, *KN1*, meristem (clusters 13, 10, 9, 3, and 5), ground tissue (clusters 1 and 12) and vasculature (part of cluster 9); *ZmHDZIV6*, epidermis (clusters 2 and 4); *ZmCLE7*, stem cells (clusters 10 and 13); *ZmWUS1*, organizing center (clusters 10 and 13); *BIF4*, spikelet pair meristem (cluster 10); *ZmCycB1/2*, G2/M phase (clusters 6 and 8); *ZmHis2b1*, S phase (clusters 0, 7, and 10); *BAF1*, adaxial meristem periphery (cluster 5 and part of cluster 2); *RA3*, meristem base (cluster 3); *ZmYAB8*, determinate lateral organ (cluster 15 and part of cluster 2); *ZmTMO5-LIKE2*, xylem (part of cluster 9); *ZmSHR1*, bundle sheath (part of cluster 9); *ZmDOF5.6*, phloem (part of cluster 9); *ZmNPF6.4-LIKE2*, cortex (cluster 1); *2G039074*, pith (cluster 12); *BD1*, meristem boundary (part of cluster 9). Color scale indicating normalized expression level.

**Figure S9:**
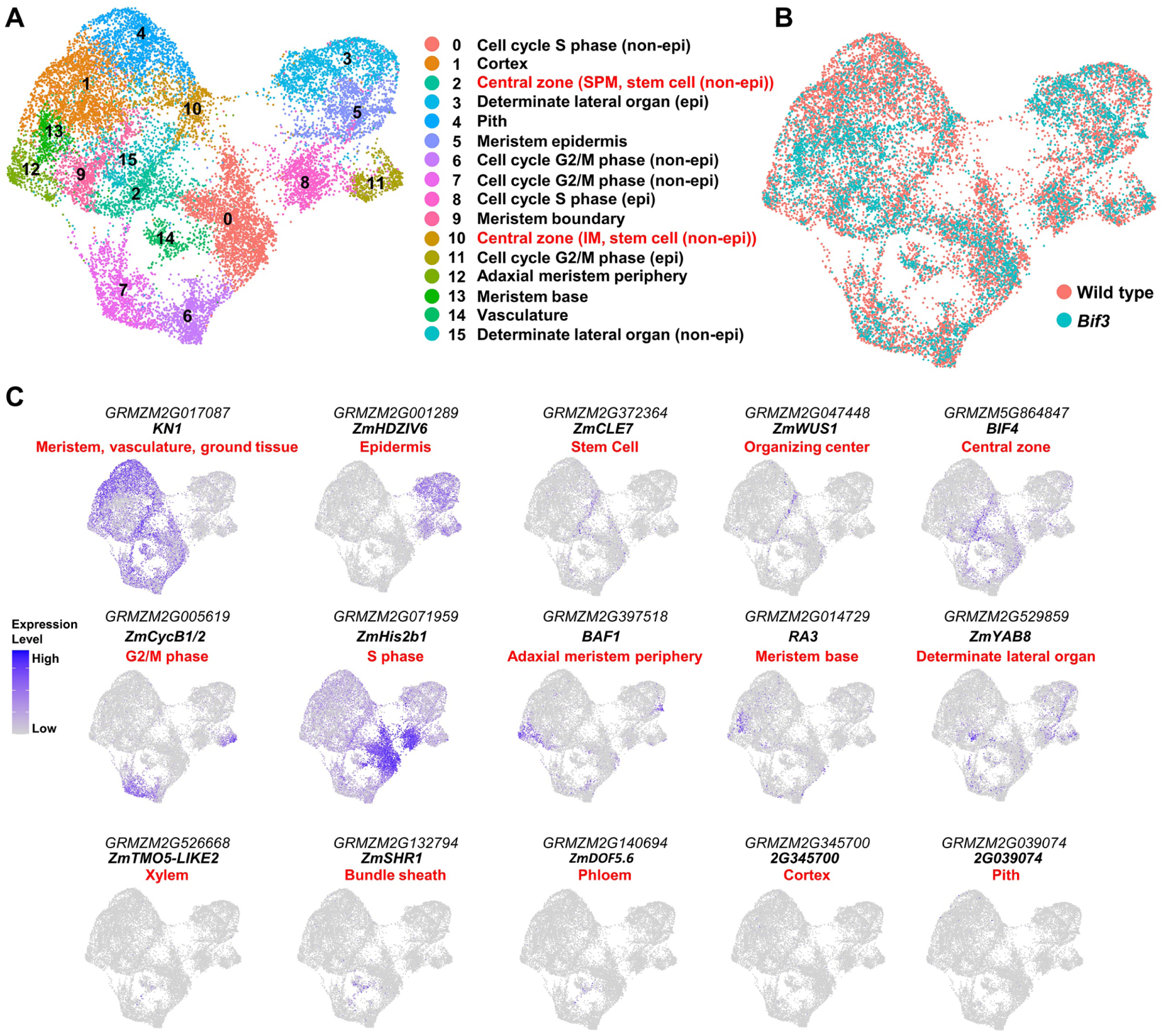
Integration of wild type and *Bif3* ear tip cell clusters. (A-B) UMAP plots showing 16 cell clusters (A) after integrating maize wild type and *Bif3* ear tip cells (B). (C) UMAP plots of marker genes predicting cluster identities, *KN1*, meristem (clusters 10, 2, 9, 13, and 12), ground tissue (clusters 1 and 4) and vasculature (cluster 14); *ZmHDZIV6*, epidermis (clusters 3 and 5); *ZmCLE7*, stem cells (clusters 10 and 2); *ZmWUS1*, organizing center (clusters 10 and 2); *BIF4*, spikelet pair meristem (cluster 2); *ZmCycB1/2*, G2/M phase (clusters 7, 6, and 11); *ZmHis2b1*, S phase (clusters 0 and 8); *BARREN STALK FASTIGIATE1* (*BAF1*), adaxial meristem periphery (clusters 12 and part of 5); *RA3*, meristem base (part of cluster 13); *ZmYAB8*, determinate lateral organ (clusters 15 and 3); *ZmTMO5-LIKE2*, xylem (cluster 14); *ZmSHR1*, bundle sheath (cluster 14); *ZmDOF5.6*, phloem (cluster 14); 2G345700, cortex (cluster 1); *2G039074*, pith (cluster4). Color scale indicates normalized expression level.

**Figure S10:**
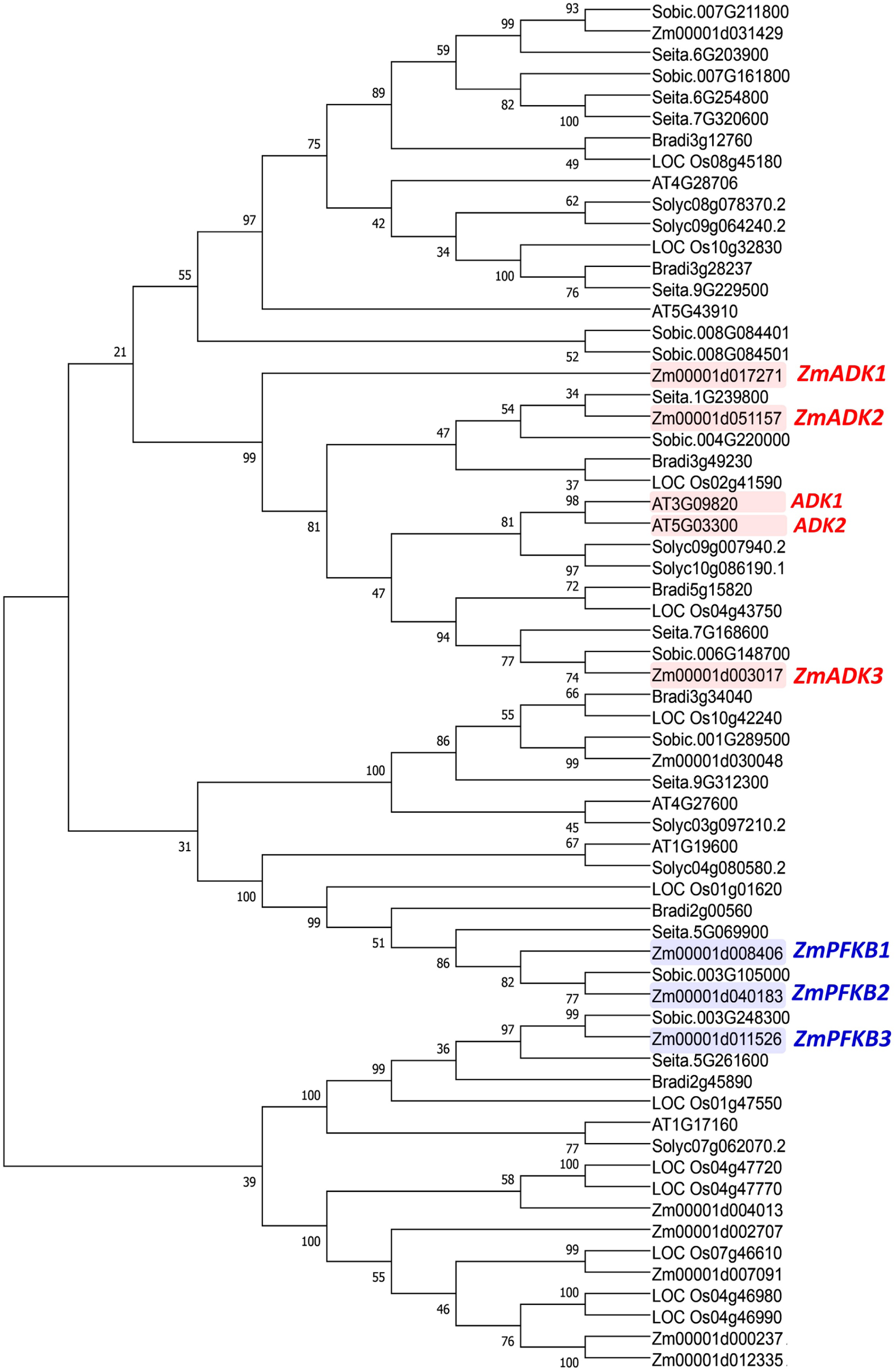
*ZmPFKB* and *ZmADK* phylogeny and expression in vegetative shoot apical meristem.

**Figure S11:**
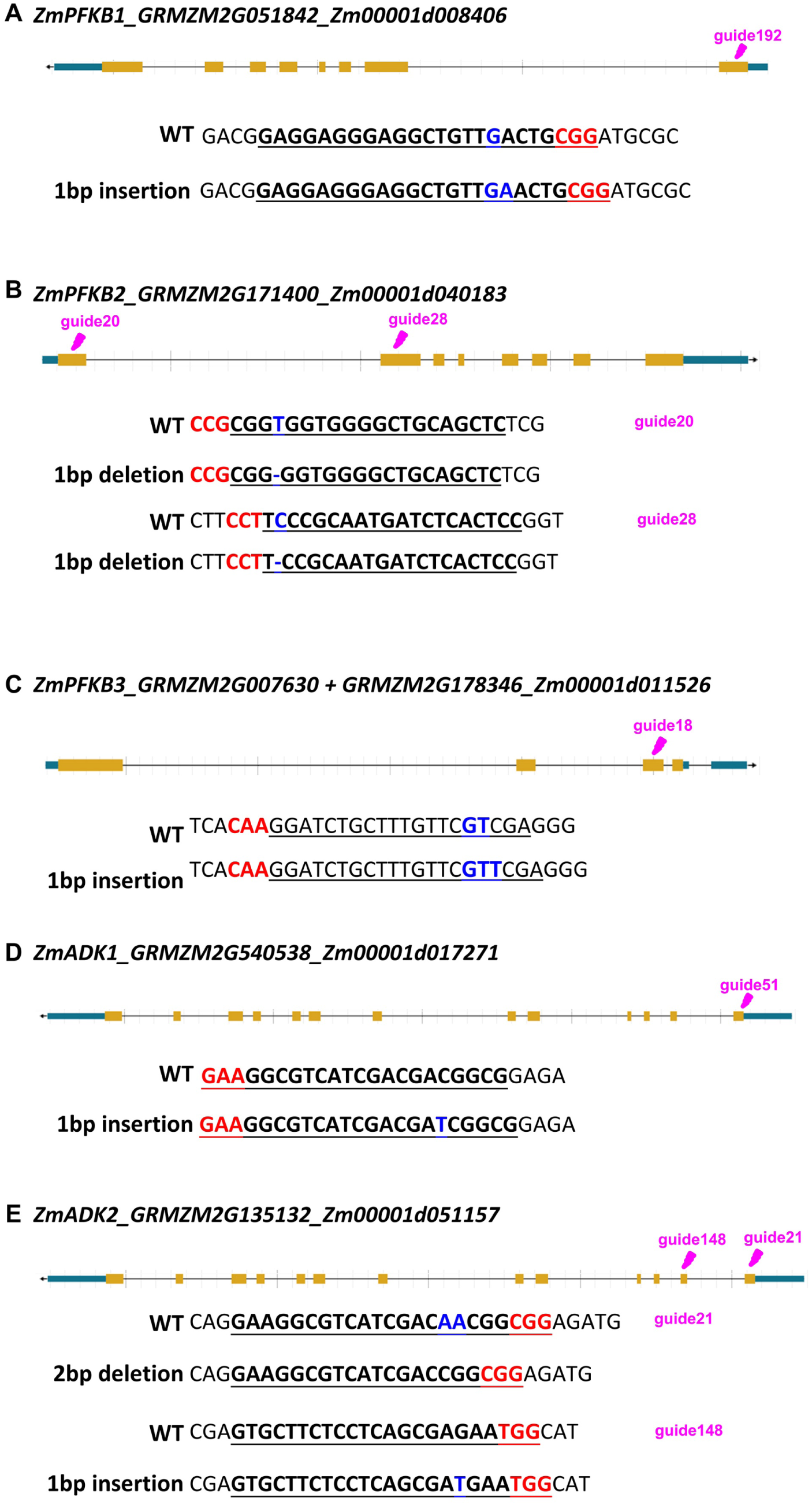
gRNA and edits for *ZmPFKBs* and *ZmADKs*. CRISPR gRNA-sequences and edits for *ZmPFKB1* (A), *ZmPFKB2* (B), *ZmPFKB3* (C), *ZmADK1* (D), and *ZmADK2* (E).

**Figure S12:**
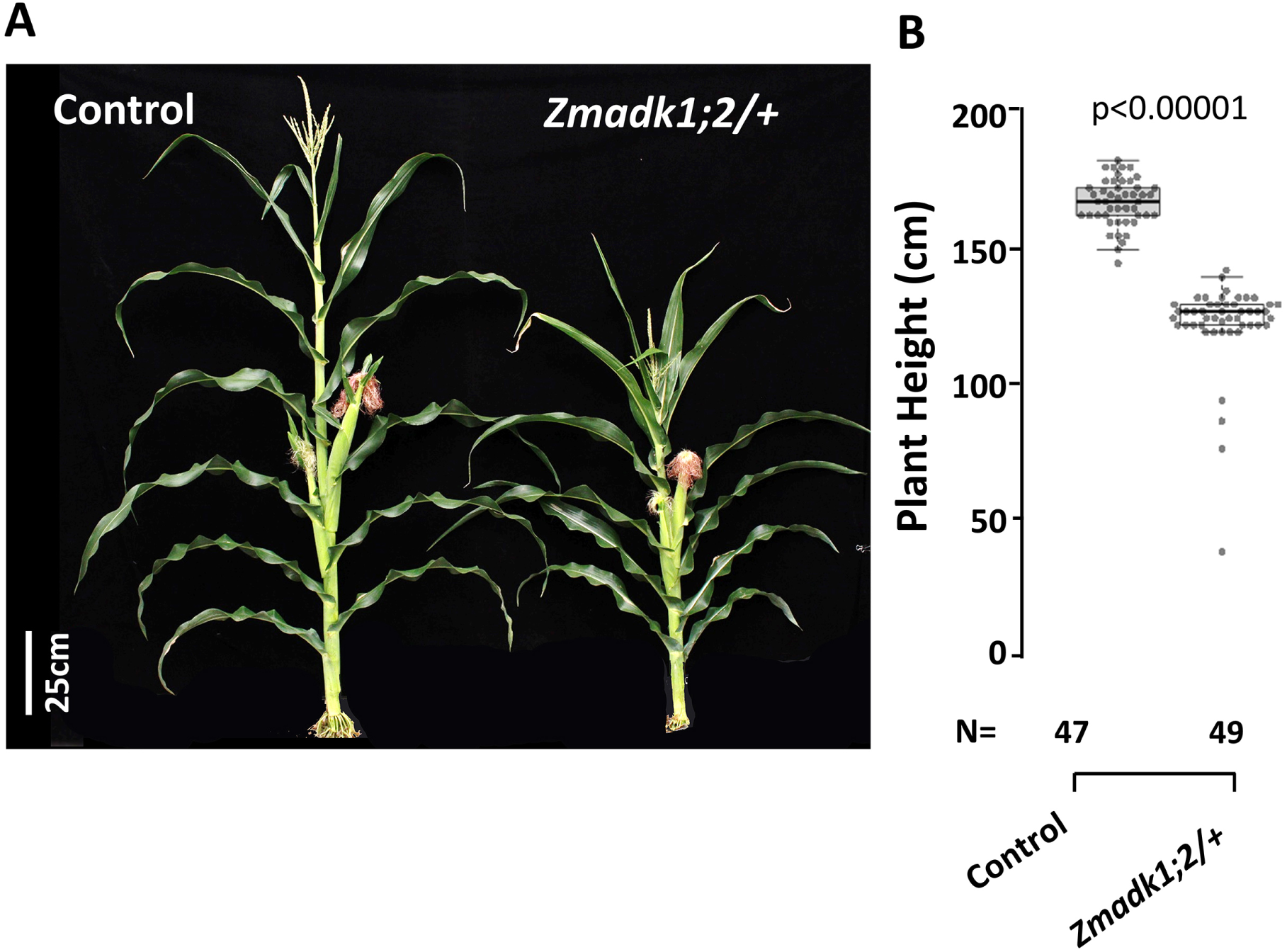
*Zmadk1;2/+* CRISPR mutants are dwarfed. *Zmadk1;2/+* CRISPR mutants have dwarf phenotype (A), with quantification shown in box plots (B) (One-Way ANOVA, p<0.00001). The horizontal line within the box represents the median.

